# Cell culture NAIL-MS allows insight into human RNA modification dynamics *in vivo*

**DOI:** 10.1101/2020.04.28.067314

**Authors:** Matthias Heiss, Felix Hagelskamp, Stefanie Kellner

## Abstract

In the last years, studies about the dynamics of RNA modifications are among the most controversially discussed. As the main reason, we have identified the unavailability of a technique which allows to follow the temporal dynamics of RNA transcripts in human cell culture. Here, we present a NAIL-MS (nucleic acid isotope labeling coupled mass spectrometry) scheme for efficient stable isotope labeling in both RNA and DNA (>95% within 7 days) in common human cell lines and growth media. Validation experiments reveal that the labeling procedure itself does neither interfere with the isotope dilution MS quantification nor with RNA modification density. We design pulse chase NAIL-MS experiments and apply the new tool to study the temporal placement of modified nucleosides in *e.g.* tRNA^Phe^ and 18S rRNA. In existing RNAs, we observe a constant loss of modified nucleosides over time which is masked by a post-transcriptional methylation mechanism and thus not detectable without NAIL-MS. During alkylation stress, NAIL-MS reveals an adaptation of tRNA modifications in new transcripts but not existing transcripts.

Overall, we present a fast and reliable stable isotope labeling strategy which allows a more detailed study of RNA modification dynamics in human cell culture. With cell culture NAIL-MS it is finally possible to study the speed of both modification and demethylation reactions inside human cells. Thus it will be possible to study the impact of external stimuli and stress on human RNA modification kinetics and processing of mature RNA.

## Introduction

Most RNAs studied to date were found to be covalently modified by dedicated enzymes in a site specific manner. In addition to the placement of RNA modifications by RNA writer enzymes, their direct removal through *e.g.* demethylation by RNA erasers was reported. In human cells, the α-ketoglutarate dependent dioxygenases ALKBH5 and/or FTO were found to catalyze the demethylation of *e.g.* (2’-O-methyl-)N6-methyladenosine (m^6^A(m)) in mRNA ^1,2^ and thus influence *e.g.* the stability and translational function of mRNA ^1,3-8^.

For human tRNAs, a similar relationship of RNA writers and erasers was observed. *E.g.* ALKBH1 demethylates 1-methyladenosine (m^1^A) and appears to be responsive to glucose starvation in some cell lines ^9^. Considering the half-life of mammalian tRNAs (∼ 100 h ^10^), a fast adaptation by removal of modified residues appears beneficial to react to changes in the cellular environment ^11^. Unfortunately, it is currently not possible to analyze the speed of both modification and demethylation reactions inside human cells. Thus it is not possible to study the impact of external stimuli and stress on human RNA modification kinetics and processing of mature RNA.

tRNA is the most extensive and chemically diverse modified RNA with ∼10-15% of all nucleosides being modified ^12^. Recent studies showed that certain modified nucleosides in specific tRNAs are only partially modified ^13,14^ and that tRNA modification abundance differs among tissues ^15,16^. This would allow for an adaptation of translation by tRNA modification as recently suggested ^17^. While the speed of tRNA amino acid charging ^18^ and tRNA transcription and half-live are known ^10^, the speed of modification processes is difficult to study. For example, tRNA^Phe^ is heavily post-transcriptionally modified and in addition one of the best studied RNAs ^19-21^. By using stable isotope labeled tRNA^Phe^ substrate and cellular extracts, the modification dynamics and hierarchy was recently solved in *S. cerevisiae* using NMR spectroscopy ^22^. Under the influence of chemical stress, *S. cerevisiae* was reported to adapt its abundance of tRNA modifications and thus influence its translation and the term stress induced tRNA reprogramming was coined ^11,23^. Similar evidence has been observed in other organisms, including mammals ^24^. In this context, the question remains by which mechanism and how fast tRNA modifications respond to external stimuli.

In contrast to tRNA, 18S rRNA is mainly modified by methylation of ribose and altogether only 2.05 % nucleosides are modified. While tRNA modifications are easily accessible for potential RNA erasers, rRNA modifications are placed in the functional regions of the ribosome ^25^. Although modified sites in rRNA have been reported to regulate translation initiation by promoting the recognition of different mRNA subsets ^26^ their inaccessibility in mature ribosomes makes them a difficult target for RNA erasers.

Current studies of RNA modifications are limited to either mass spectrometric analysis ^16^ or sequencing ^27,28^. Both techniques provide information on the modification status at the time point of sample harvest and give no details on the mechanisms of RNA modification adaptation. To overcome this limitation, we have recently developed NAIL-MS (nucleic acid isotope labeling coupled mass spectrometry) in bacteria ^29,30^ and yeast ^31^, which reveals the dynamics of RNA modification processes. The technique is based on metabolic stable isotope labeling of RNA using simple nutrients with *e.g.* carbon-13, nitrogen-15 or sulfur-34. By combining differentially labeled media in a pulse chase set-up, we recently succeeded to observe tRNA demethylation through AlkB in *E. coli in vivo*. Currently, NAIL-MS studies are not available for human cell lines as a monoisotopic labeling of all four canonical nucleosides is highly complex and thus not available.

Here, we report a fast and reliable method for monoisotopic stable isotope labeling in both RNA and DNA (>95% within 7 days) in common human cell lines and growth media. We apply the cell culture NAIL-MS method and reveal the dynamics of human tRNA and 18S rRNA modifications in depths unreachable by any other tool for RNA modification analysis. Furthermore, we resolve the mechanism of stress induced tRNA modification reprogramming in the presence of methylation stress. With cell culture NAIL-MS it is finally possible to study the speed of both modification and demethylation reactions inside human cells. Thus it will be possible to study the impact of external stimuli and stress on human RNA modification kinetics and processing of mature RNA.

## Results

### Absolute quantification of human tRNA^Phe^ modifications

tRNA^Phe^ is heavily post-transcriptionally modified and in addition one of the best studied RNAs ^19-22^. Thus it is an ideal model to study the temporal dynamics of its modifications. In a first step, we purified tRNA^Phe^_GAA_ from HEK 293 cells using a complementary DNA probe ^13^. We used our established isotope dilution LC-MS/MS analysis for absolute quantification of modified nucleosides and plotted the modification profile in Figure 1 ^16^. For pseudouridine (Ψ), dihydrouridine (D), 2-dimethylguanosine (m^22^G) and 2’-O-methylguanosine (Gm) our experimental data matches the expected values and we see full modification ^32^. The abundance of 1-methyladenosine (m^1^A position 14 and 58), 7-methylguanosine (m^7^G), 5-methyluridine (m^5^U or rT) and 2’-O-methylcytidine (Cm) is lower compared to the literature, presumably due to partial modification of the respective sites. Partial modification has been suggested to play a role in stress induced reprogramming of tRNA modifications ^17^. The abundance of 5-methylcytosine (m^5^C) is slightly higher than expected and can be explained by the additional methylation of C48 by NSUN2 ^33^.

**Fig. 1:**
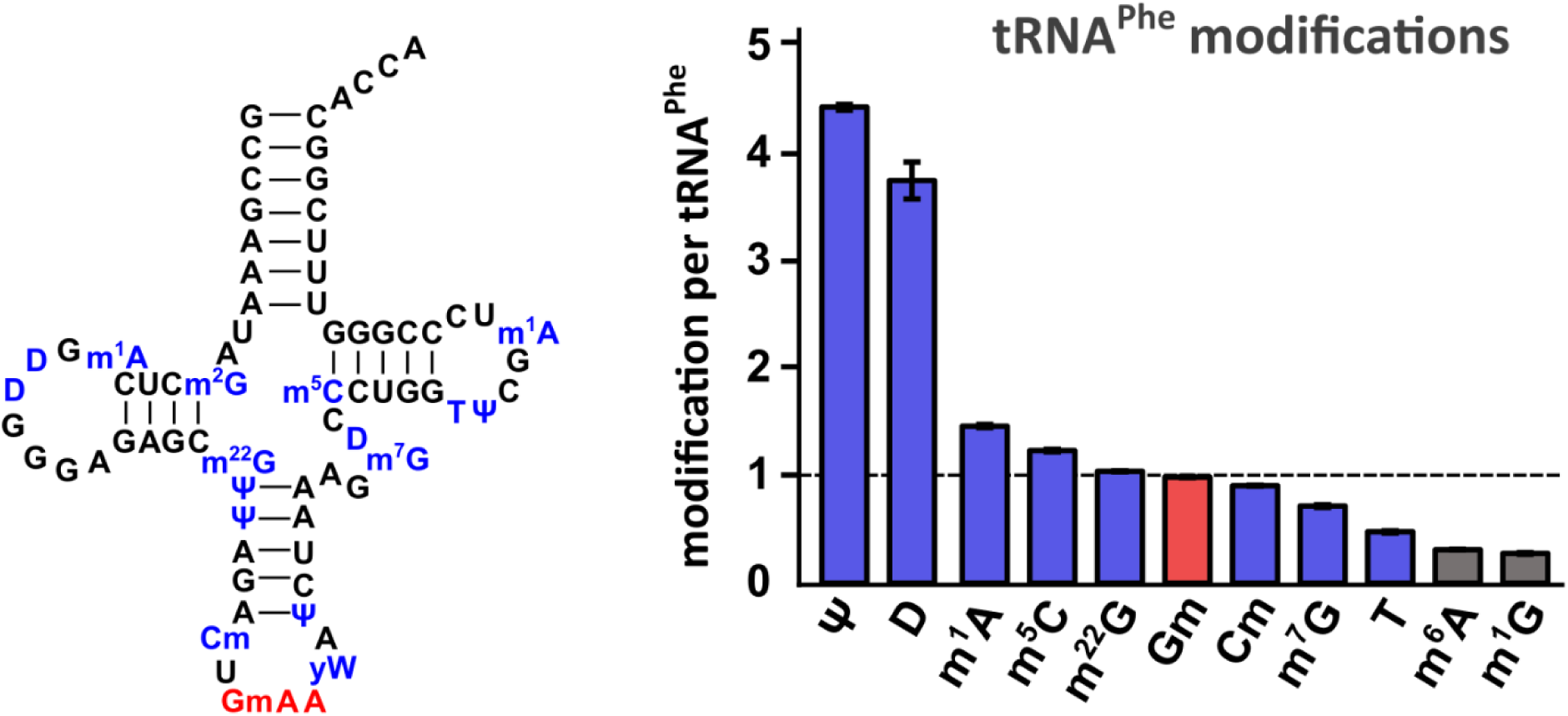
Absolute quantification of human tRNA^Phe^_GAA_ modifications. The tRNA cloverleaf on the left shows the expected sequence of human tRNA^Phe^_GAA_ including reported modifications ^32^. Right: Absolute quantification of purified tRNA^Phe^_GAA_ from HEK 293 cells done by LC-MS/MS. All experiments are from n = 3 biol. replicates and error bars reflect standard deviation.

Although 1-methylguanosine (m^1^G) is not reported in tRNA^Phe^_GAA_, we found around 0.3 m^1^G per tRNA. This observation can be explained by the fact that m^1^G is a precursor during the biosynthesis of wybutosine (yW), a hypermodified nucleoside reported at position 37 of tRNA^Phe^_GAA_ ^34,35^. Due to the unavailability of a synthetic standard, yW could not be quantified in this study. Additionally, we also quantified the abundance of other modified nucleosides (Table S1). We found around 0.3 6-methyladenosine (m^6^A) per tRNA, potentially caused by intracellular dimroth rearrangement of m^1^A ^36^. In addition, we found 0.063 inosine (I) and 0.026 1-methylinosine (m^1^I) per tRNA^Phe^. These are most likely artefacts from A and m^1^A deamination. All other modified nucleosides, were found with an abundance of less than 1.6% (*e.g.* 0.016 N^6^-threonylcarbamoyladenosine (t^6^A) per tRNA) which indicates a high purity of isolated tRNA^Phe^.

Overall the detected quantities of modified nucleosides from purified tRNA^Phe^_GAA_ are in accordance with the reported values and thus it is a suitable model to study the temporal placement of modified nucleosides.

### Stable isotope labeling of RNA in human cell culture

For this purpose, a method is needed which allows the discrimination of mature RNA from new transcripts. NAIL-MS (nucleic acid isotope labeling coupled mass spectrometry) relies on the metabolic incorporation of stable isotope labeled nutrients into RNA and allows the distinction of original RNA and new RNA within a pulse chase experiment. With this tool, we studied the temporal placement of modified nucleosides in *S. cerevisiae* total tRNA ^31^ and the demethylation during tRNA repair in *E. coli* ^37^. Both organisms are rather simple and they can be grown in minimal media with controlled availability of stable isotope labeled nutrients.

In contrast, human cell culture medium is highly complex and requires the addition of fetal bovine serum (FBS). FBS is a natural product of undefined composition and variable concentration of metabolites. Thus a complete and monoisotopic labeling of nucleosides and even nucleobases for a pulse chase NAIL-MS assay is challenging.

From our experience, the target isotopologue of a nucleoside must be at least 3 u heavier compared to the naturally occurring nucleoside to avoid false positive results by the detection of the natural carbon-13 signals.

*De novo* synthesis of nucleosides utilizes several amino acids such as glutamine or aspartic acid (Figure S1A and S1B) ^38^. Hence, we supplemented the growth media with stable isotope labeled glutamine. After 5 days (2 passages), we observed the expected stable isotope labeling of RNA (Figure S1C). Cytidine, guanosine and adenosine got a mass increase of +2 whereas uridine just increased by +1. Due to the overlap with naturally occurring (^13^C)-isotopologues, this mass increase was not sufficient for our planned experiments.

As recently described, it is possible to use glucose-free growth medium and supplement with ^13^C_6_-glucose ^37^. The feeding with ^13^C_6_-glucose leads to the formation of nucleosides with a variable number of (^13^C) per nucleoside (Figure S1C). During method development, we utilized the non-monoisotopic nature of ^13^C_6_-glucose labeling to test the incorporation efficiency of various unlabeled metabolites. Addition of aspartate and pyruvate did not allow the envisioned monoisotopic labeling (Figure S2). The addition of the nucleobases adenine and uracil resulted in ribose labeled purines but undefined labeled pyrimidines. This indicates a direct usage of adenine from the medium which is then enzymatically connected with ^13^C_5_-ribose followed by further processing to guanosine and the respective triphosphates (Figure S1B). RNA supplemented with the nucleosides adenosine and uridine showed undefined labeled purines and only unlabeled pyrimidines (Figure S3). This indicates that uridine is taken up by the cells and immediately utilized for cytidine and RNA synthesis (Figure S1A). In summary, our data indicates that the addition of adenine and uridine blocks *de novo* purine and pyrimidine synthesis (Figure S1A and S1B) and ^13^C_6_-glucose medium is not necessary for our labeling strategy as unlabeled nucleosides remain visible in the mass spectra (Figures S3). Concentration optimization of both compounds revealed that final concentrations of 0.1 mM adenine and 0.2 mM uridine in the ^13^C_6_-glucose medium are needed to suppress signals from de novo synthesized nucleosides (Figure S4).

Instead, we used ^15^N_5_-adenine and ^13^C_5_,^15^N_2_-uridine (Figure 2a) in medium with unlabeled glucose. The high resolution mass spectra of the resulting RNA nucleosides showed the desired labeling for >95% of all canonical nucleosides after 7 days (Figure 2b). A +7 mass increase is observed for cytidine and uridine and a +5 and +4 mass increase for adenosine and guanosine, respectively. By using dialyzed FBS, the signal of unlabeled adenosine could be further reduced in comparison to normal FBS (Figure S5). Similarly, DNA nucleosides become stable isotope labeled (Figure S6). With these metabolites, a pulse chase NAIL-MS study is possible in human cell culture. To this end, we achieve excellent labeling in HEK 293, HAP and HeLa cell lines using supplemented DMEM RPMI or IMDM medium (Figure S7).

**Fig. 2:**
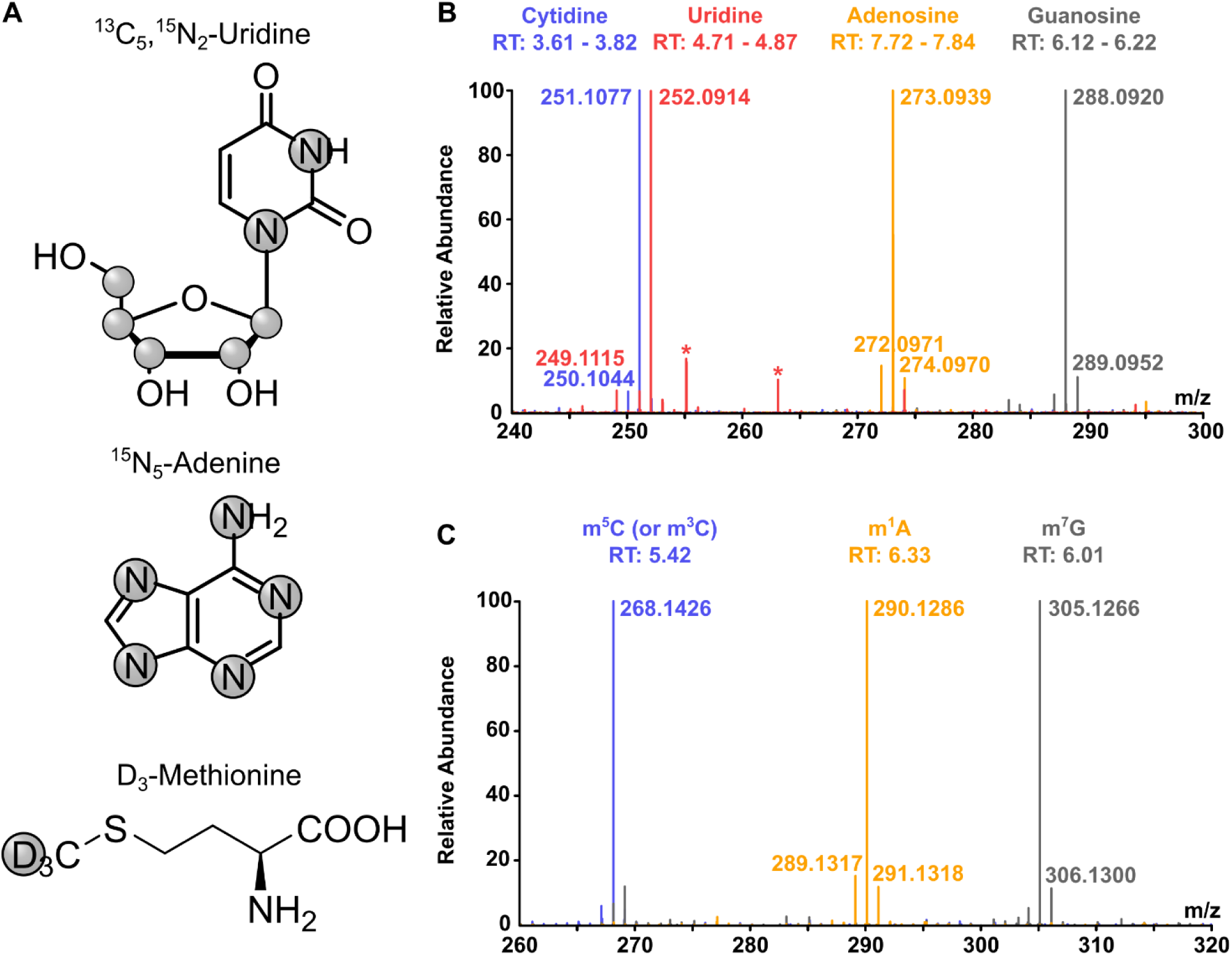
High resolution mass spectra of stable isotope labeled nucleosides from cell culture. **a**, Labeling of compounds used for stable isotope labeling in cell culture. Grey circles indicate the positions of isotopes (^13^C, ^15^N or ^2^H/D). **b**, Merged high resolution mass spectra of the 4 canonical nucleosides of total tRNA after labeling of HEK 293 cells with compounds shown in a for 7 days. Background signals are marked with asterisks. **c**, Merged high resolution mass spectra of three exemplary modifications (m^5^C, m^7^G and m^1^A) in total tRNA after stable isotope labeling of HEK 293 cells.

In mouse embryonic stem cells (mESC), the addition of ^15^N_5_-adenine and ^13^C_5_,^15^N_2_-uridine leads to non-monoisotopic labeling. Here, the labeling efficiency is improved from 35% to 70% by the usage of dialyzed FBS (Figure S8).

In HEK 293 cells, the signals of new tRNA transcripts became detectable and quantifiable after 120 minutes of labeling (Figure S9/S10).

Most modified nucleosides in RNA carry one or more methylations. To follow the fate of these methylated nucleosides in the context of RNA maturation and methylation damage response, we used CD_3_-labeled methionine. Methionine is the precursor amino acid of S-adenosylmethionine (SAM) which in turn is cofactor of most RNA methyltransferases. In the presence of CD_3_-methionine, methylated nucleosides get a mass increase of +3 and can thus be distinguished from nucleosides modified in the presence of unlabeled methionine. High resolution mass spectra of fully labeled m^5^C, m^7^G and m^1^A are exemplarily shown in (Figure 2c). In order to achieve complete labeling of methyl-groups methionine depleted medium has to be used. We chose DMEM D0422 (from Sigma-Aldrich) which lacks glutamine, cystine and methionine (Figure S11). Neither cell shape nor growth speed were influenced by the labeling and both were comparable to standard DMEM (*e.g.* D6546, from Sigma-Aldrich) (Figure S12).

The combination of nucleoside and methyl-group labeling allows the design of elegant pulse chase studies to follow the fate of RNA in human cells.

### Validation of human cell culture NAIL-MS

After finding a suitable way for monoisotopic labeling of RNA in human cells, we wanted to rule out the possibility that the labeling itself impacts the abundance of RNA modifications. For this purpose, cells were grown in labeled or unlabeled media for 7 days. Both media contained adenine, uridine and methionine as either unlabeled or labeled nutrients. Cells were harvested with TRI reagent and split into two aliquots. One aliquot (2/3 Vol) was used for immediate RNA isolation and purification, while the remaining aliquot of the labeled and unlabeled cells were mixed and RNA was co-isolated and co-purified (Figure 3a and Figure S13). The total tRNA was enzymatically digested to nucleosides and their abundance determined by isotope dilution mass spectrometry ^16^. In the aliquot from unlabeled samples, only unlabeled nucleosides were detectable, while the aliquot of the labeled cells showed mainly signals (>98%) for labeled nucleosides. As expected from the mixed sample, we detected unlabeled and labeled isotopologues of all canonicals in equivalent amounts (Figure 3b). Next, we quantified the abundance of modified nucleosides. For normalization, unlabeled modifications were referenced to unlabeled canonicals and labeled modifications were referenced to labeled canonicals. The calculated quantities of modified nucleosides present in tRNA^Phe^ are plotted for the unlabeled against labeled tRNA in Figure 3c. This validation revealed that the quantities of modified nucleosides are independent of the media and that the labeling procedure itself does not interfere with the isotope dilution MS quantification. The deviation from the expected values is the error of this NAIL-MS experiment and the limitation to detect differences in a biological setup (also see Figure S14). *E.g.* In total tRNA, 2’-O-methyluridine (Um) has the largest error as its abundance deviates 1.6 fold in labeled and unlabeled media.

**Fig. 3:**
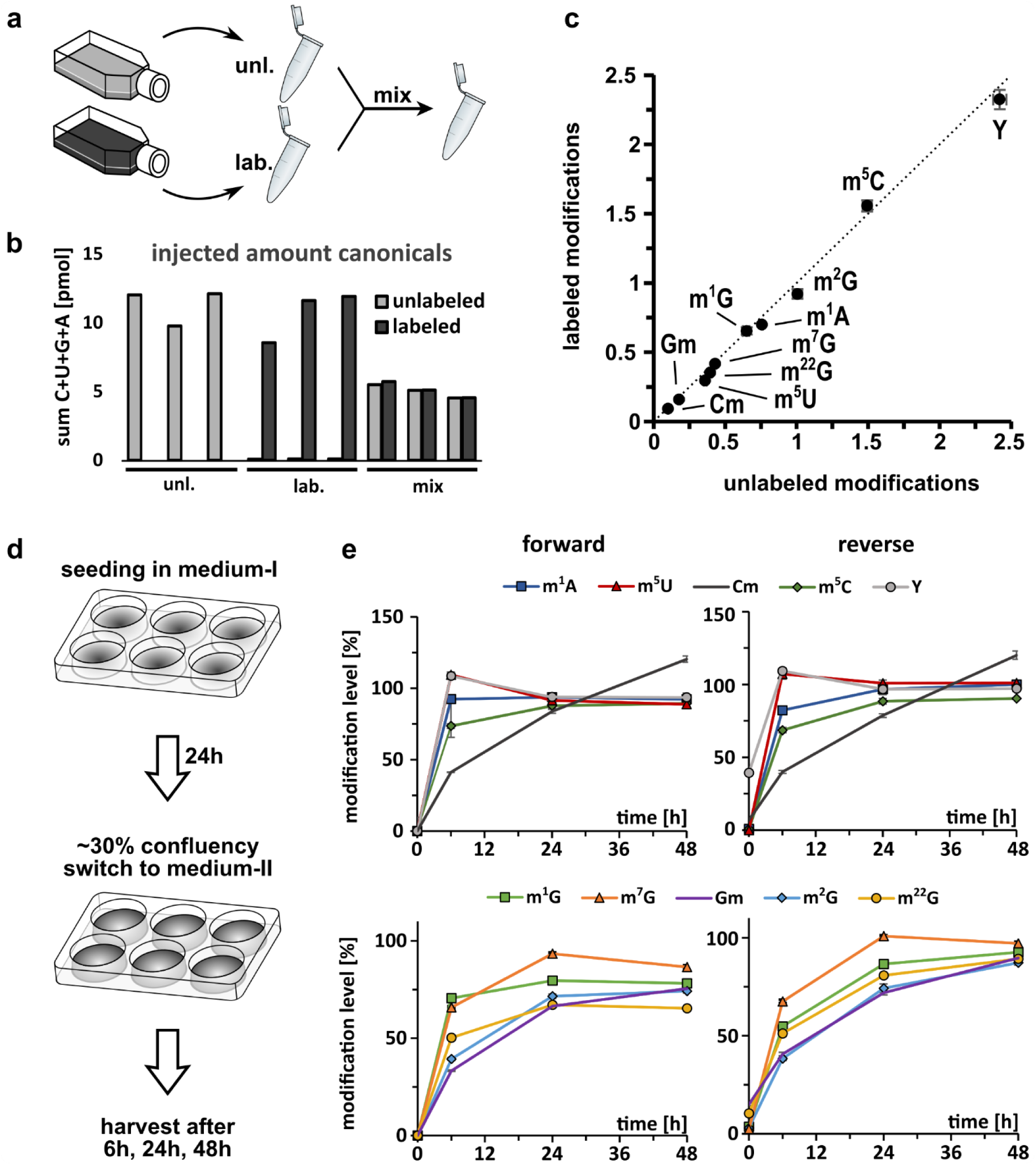
Validation of cell culture NAIL-MS. **a**, Cells were grown in unlabeled or fully labeled media for 7 days. Upon harvesting one aliquot was mixed prior to processing (mix). Total tRNA was purified and all samples were analyzed by LC-MS/MS. **b**, Summed amount of canonical nucleosides (C+U+G+A) detected by LC-MS/MS for unlabeled and labeled isotopomers. The bars show single replicates of three unlabeled, three labeled and three mixed aliquots **c**, Abundance of labeled modifications plotted against the abundance of unlabeled modifications in the mix samples. The dotted line indicates the location of the expected values as a visual guide. **d**, Experimental setup of time course study to investigate temporal placement of RNA modifications. The experiment was done forward (start with unlabeled, change to labeled medium) and reverse *(vice versa*). **e**, Results of time course study. Plotted on the y-axis is the abundance of modification in new transcripts normalized to the abundance before experiment initiation (T = 0). Note: In the reverse experiment, minor signals of unlabeled nucleosides are present at T=0 and thus the starting value is sometimes larger than 0%. All experiments were done with purified total tRNA and are from n = 3 biol. replicates. Symbols reflect the mean and error bars reflect standard deviation.

The promising results from the validation experiments allowed the design of pulse chase experiments. Such experiments start with cells seeded in medium-I and upon experiment initiation, the medium is exchanged to medium-II with different isotopically labeled nutrients. The concept is shown in Figure 3d. To rule out possible differences in the results in dependence of the starting medium, we designed a brief validation experiment. In the forward experiment, cells are seeded in unlabeled medium and switched to labeled medium while the reverse experiment starts in labeled medium (after a 7 day labeling period) before switching to unlabeled.

For analysis of modified nucleoside quantities, we harvested the cells and extracted total tRNA after switching to medium-II (time points 0, 6, 24 and 48 hours). To assess the suitability of the method for temporal placement of modified nucleosides into the total tRNA, we focused on the abundance of new modified nucleosides in the newly transcribed tRNA. For direct comparison, the ratio of found (6, 24, 48 h – new transcripts) and expected (0 h – original transcripts) modified nucleoside quantity was formed and plotted over time. As expected, we observed the incorporation of modified nucleosides into the new tRNA after medium exchange. While the timing of the tRNA modification process was comparable in the reverse and forward experiment, the start values were obscured in the reverse experiment due to low, but detectable signals of unlabeled nucleosides. For this and economic considerations, we decided to perform forward pulse chase experiments in the future to avoid the excessive use of labeled medium.

### Temporal placement of modified nucleosides in RNA

From a biological perspective, we observed that most modified nucleosides reach their final abundance (100 % compared to the starting point) within 48 h (Figure 3e). Some modified nucleosides, such as m^1^A, m^5^C, Ψ and m^5^U, are already > 90 % after 6 h which indicates a fast incorporation after transcription. These modified nucleosides are located in the structure-stabilizing positions of the tRNA’s D-and TΨC-loops and thus a fast and reliable modification is to be expected ^39^. m^7^G is also involved in structure stabilization ^40^ and yet, this methylation is placed rather slowly in total tRNA. Other modified nucleosides such as Cm, Gm, and the base-methylated G derivatives (m^1^G, m^2^G and m^22^G) are incorporated more slowly and the final modification density is not reached within 48 h.

While the modified nucleosides of total tRNA are placed by various enzymes at various positions, we were interested to observe the modification process of defined enzymes in a defined substrate. For this purpose, we performed a pulse chase experiment and purified tRNA^Phe^_GAA_ after 0, 2, 4, 6, 24 and 48 hours. The abundance of modified nucleosides in new tRNA transcripts is shown in Figure 4. We observe an immediate high abundance of Ψ, which argues towards an immediate isomerization of *e.g.* U55 to Ψ55 as observed in yeast ^22^. In fact, we observe 1.5 fold more Ψ in the early lifetime of tRNA^Phe^ _GAA_ as is expected from mature tRNA^Phe^_GAA_ (Figure 1). At these early time points, the abundance of new tRNA^Phe^ _GAA_ transcripts is low and thus the MS signal intensity is close to the lower limit of quantification (LLOQ). Uridine and its modifications have a low ionization efficiency and thus a higher LLOQ compared to other modified nucleosides. Thus biological interpretation of Ψ and m^5^U (Figure S15) quantities must be conducted carefully. D is not included in this analysis, due to its artificial addition to the samples through the deaminase inhibitor tetrahydrouridine (which was omitted for analysis in Figure 1 and thus allowed quantification of D). While m^7^G is the next modified nucleoside placed in yeast tRNA^Phe^, our data hints towards a fast incorporation of m^5^C followed by m^1^A and finally m^7^G. Here, the dynamic placement of modifications in the TΨC-loop seems to be slightly different between yeast and human. The slow incorporation of m^2^G in the D-loop is in accordance with the reports from yeast. In the anticodon-loop (ac-loop), we observe a rather slow formation of Gm and Cm. These modified nucleosides are not involved in structure stabilization but codon-anticodon binding ^41,42^ and protein translation. Our data implies that structure stabilization by modified nucleosides is a key necessity and must thus happen early on, while ac-loop modifications are not immediately needed and are potentially placed on-demand. One exception is the formation of wybutosine (yW). Its precursor modification m^1^G is immediately incorporated into tRNA^Phe^ before its abundance drops at later time points, presumably due to its further processing into yW.

**Fig. 4:**
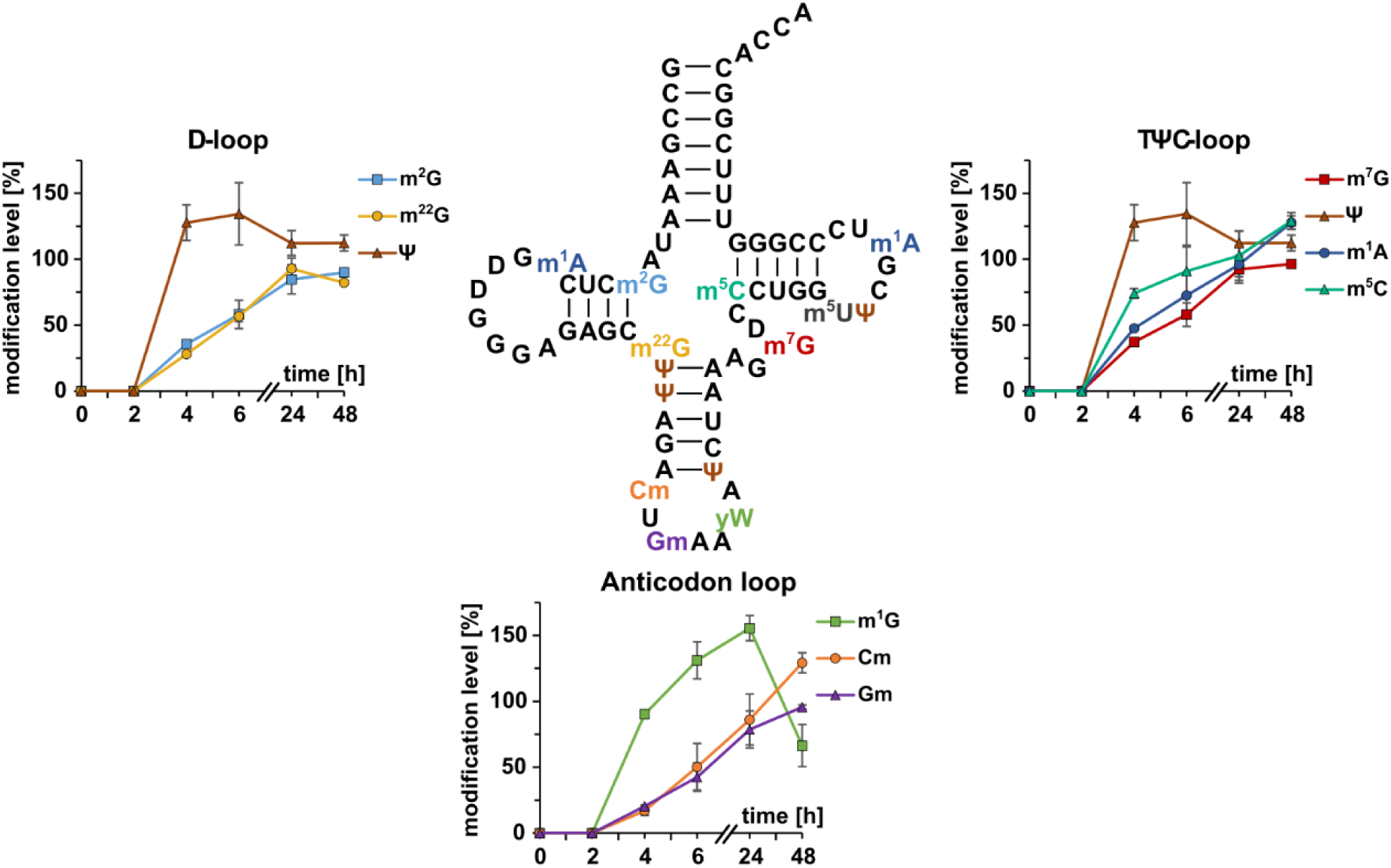
Temporal placement of modified nucleosides in tRNA^Phe^_GAA_. Cells were grown in unlabeled DMEM D0422 (supplemented with unlabeled uridine and adenine) for 7 days. At T = 0 the medium was exchanged to DMEM D0422 supplemented with labeled uridine and adenine. Cells were harvested after set time points. tRNA^Phe^ was purified and analyzed by LC-MS/MS. Modifications are plotted next to their location in the D-, TΨC-or anticodon loop. Plotted on the y-axis is the abundance of modification in new transcripts normalized to the respective nucleoside originating from unlabeled medium before experiment initiation (T = 0). The experiment was done in n = 3 biol. replicates for time points 2, 4 and 48 h and in n = 6 biol. replicates for time points 6 and 24 h. Symbols reflect mean and error bars reflect standard deviation.

### Dynamics of tRNA and 18S rRNA modifications

With the design of our pulse chase NAIL-MS assay, we can observe RNA maturation processes by quantifying the abundance of modified nucleosides in new transcripts. In addition, we can follow the fate of original RNA (unlabeled nucleosides in forward experiment) and observe methylation or demethylation events.

In Figure 5a, we plotted the abundance of exemplary modified nucleosides from original total tRNA, which were present before the medium exchange. Other modified nucleosides are shown in Figure S16. Similar to our initial observations in *S. cerevisiae* ^31^, we observed a constant loss of modified nucleosides from original tRNAs. In the common, unlabeled analysis of modified nucleosides, the decrease in modification density from original tRNA is not visible as it is masked by the addition of new methyl marks to original tRNAs at early time points (post-methylation) and by quickly modified new transcripts at later time points (ratio original/new transcripts in Figure S16/17). Here, the post-methylation reaction is captured by the CD_3_-methionine added in the chase phase (medium II) and is termed “methyl” in Figure 5a. Intriguingly, the extent of post-methylation depends on the modified nucleoside. For m^7^G and Cm, it is more pronounced compared to m^1^A. Interestingly, many modified nucleosides which are placed almost immediately after transcription, show low amounts of post-methylation while those with a delayed incorporation showed substantial post-methylation. Similar to tRNA, ribosomal RNA nucleosides are modified, mainly at locations close to the functional region of the ribosome ^25^. From yeast studies, it is known that most ribose rRNA modifications are inserted immediately or even co-transcriptionally ^43^. For Ψ and other base modifications, the time point of placement during rRNA maturation is yet unknown. In a forward pulse-chase experiment, we have isolated the 18S rRNA and quantified the abundance of the original and new modified nucleosides. As expected from yeast, ribose methylations appear early on in the new 18S rRNA transcripts. Intriguingly, m^6^A and Ψ are inserted as fast or even faster. This indicates an immediate placement after transcription which is in agreement with their inaccessibility at later stages of ribosome biogenesis (Figure S18).

**Fig. 5:**
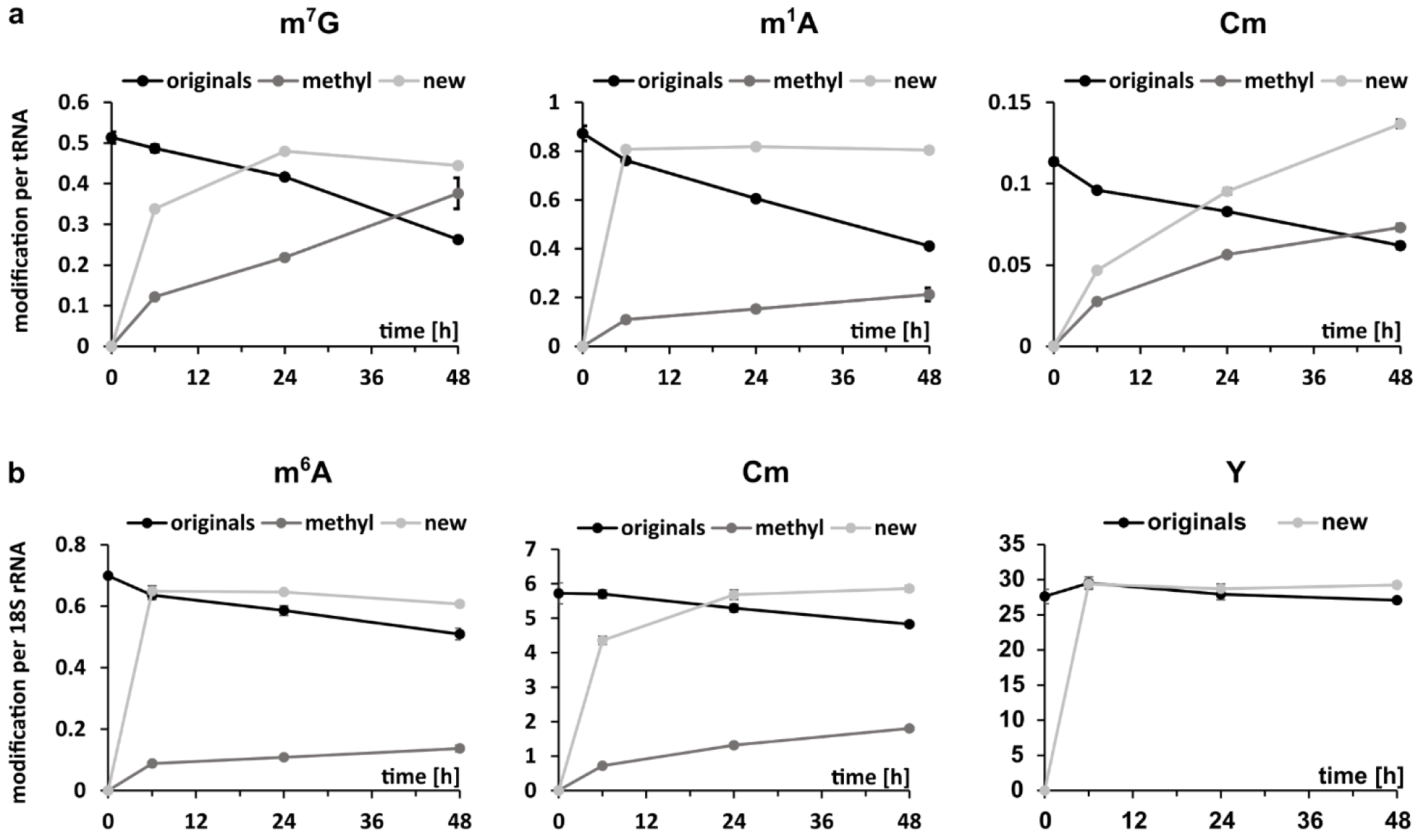
Dynamics of modified nucleosides in total tRNA (a) and 18S rRNA (b). Original nucleosides (originals, black line) existed before experiment initiation. Post-methylated nucleosides (methyl, dark grey line) are modifications arising from the methylation of original RNA after experiment initiation. New nucleosides (new, light grey line) show the incorporation of modification into new transcripts. Data points reflect the mean and standard deviations of n = 3 biol. replicates.

### Impact of methylation stress on tRNA modification processes

We have recently applied NAIL-MS to profile bacterial tRNA damage by methylating agents ^29^ and described the repair mechanisms *in vivo* ^37^. With the goal to study the stress response in human cells, we determined the effect of methyl methanesulfonate (MMS) on growth of HEK 293 cells (Figure S19). In these experiments, we observed a strong influence of trypsinization on cell survival, which we avoided in later experiments.

Until now, it was not possible to study the extent of m^1^A and m^7^G damage formation in human RNA due to the presence of enzymatically placed m^1^A and m^7^G. With our cell culture labeling scheme we succeeded to implement a methylome discrimination assay and determine the absolute abundance of these major RNA damages. For this purpose, cells were grown in CD_3_-methionine supplemented medium for 7 days before addition of 1 mM MMS. While enzymatically placed methylations are CD_3_-labeled, MMS damaged sites are CH_3_-labeled and thus easily distinguishable from the enzymatic sites by mass spectrometry. To enable the tracing of the damaged tRNAs without interference of new transcripts, we included a switch to fully labeled media (^13^C/^15^N-nucleoside + CD_3_) for labeling of new transcripts. The time line and concept is given in Figure 6a. Samples of MMS and MOCK treated cells were taken before and after 1 h of MMS exposure and up to 6 h after removal of MMS, where cells were left to recover from the treatment.

**Fig. 6:**
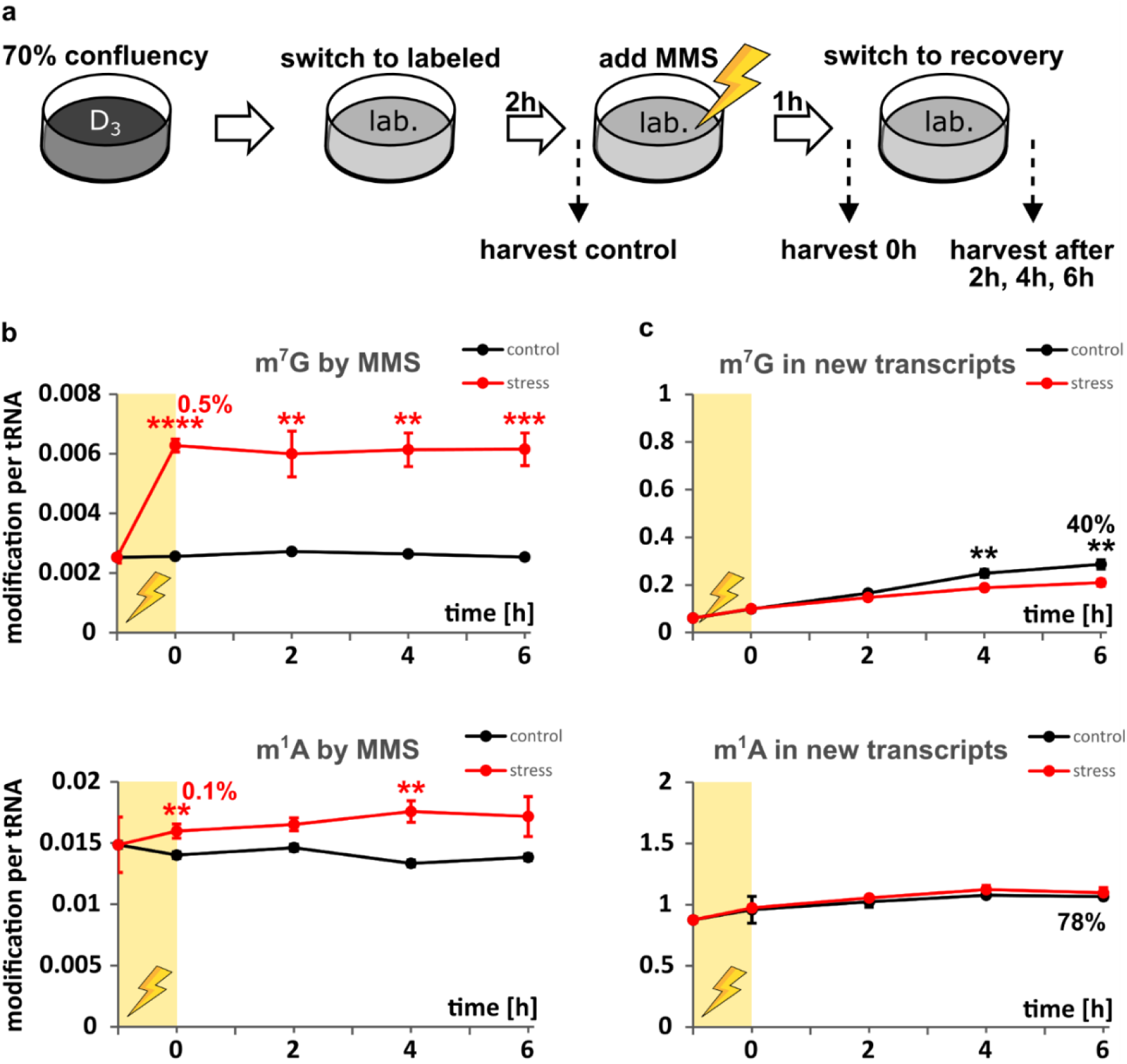
Effect of methylation stress on tRNA modifications. **a**, 70% confluent CD_3_-methionine labeled cells were incubated with fully labeled media for 2 h before the LD_50_ dose of methyl methanesulfonate (MMS, yellow shaded area) was added. Control samples were treated the same with PBS instead of MMS. After 1 h the stress (or control) media was replaced by fresh labeled media. After set time points, cells were harvested and tRNA^Phe^_GAA_ was purified and subjected to LC-MS/MS analysis. **b**, Unlabeled modifications were referenced to unlabeled canonicals to calculate the amount of modifications arising from direct methylation damage by MMS. The numbers at time point 0 give the percentage of damaged nucleoside referenced to the naturally occurring amount of the respective modification. **c**, Labeled modifications were referenced to labeled canonicals to calculate the amount of modification in new tRNA transcripts. The numbers at time point 6 give the percentage of modification amount in the control sample referenced to the naturally occurring amount of the respective modification. All experiments are from n = 3 biol. replicates and error bars reflect standard deviation. P-values from student t-test (equal distribution, two-sided): * p < 0.05, ** p < 0.01, ***p < 0.001 and ****p < 0.0001.

From these samples, we purified tRNA^Phe^_GAA_ and quantified the abundance of canonical and modified nucleosides. By comparison of canonical nucleosides, we could observe a higher ratio of new transcripts over original transcripts in the unstressed samples compared to the stressed samples (Figure S20). This is to be expected as stressed cells stop growing and thus less transcription and translation are needed. In addition, the prolonged abundance of original tRNA suggests that methylation stress does not lead to extensive degradation of tRNAs.

The quantification of methylated nucleosides derived from direct MMS methylation, indeed showed formation of the known damage products m^7^G and potentially m^1^A. In comparison to the natural abundance of these modified nucleosides (∼ 0.5 m^7^G and 1 m^1^A per tRNA^Phe^_GAA_), the damage accounts for less than 1 % of these methylated nucleosides (Figure 6b). In other words, only 1 out of around 200 tRNA molecules gets an additional m^7^G by MMS damage. For m^1^A the damage is found in 1 out of 1000 tRNAs (0.1 %). No other reported MMS damage products were detected in human tRNA^Phe^.

While RNA methylation damage repair was observed in *E. coli*, using a similar NAIL-MS approach, no demethylation was detectable in the human cell line. Even 6 hours after removal of the methylating agent, the abundance of directly methylated m^7^G and m^1^A stayed unchanged in the original transcripts. This observation indicates that human cells either have a highly expressed and fast acting RNA demethylase for RNA damage repair, or the abundance of damaged nucleosides is below a threshold limit to trigger repair or there is no RNA demethylase for damage repair in human cells. We next asked the question, whether human cells react to methylation stress by adaptation of tRNA modifications. This adaptation can be mechanistically achieved by addition or removal of modified nucleosides to original tRNAs, by delayed modification of new tRNAs or a combination of both. For methylation stress, we did not see a difference in modified nucleoside abundance in original tRNA compared to the unstressed control (Figure S21). (Note: In this NAIL-MS study, the supplemented methionine was CD_3_-labeled in both the pulse and the chase phase. Thus, it is not possible to observe the decrease of enzymatically placed modifications in original tRNA over time as shown in Figure 5).

Finally, we studied the abundance of modified nucleosides in new tRNA transcripts in dependence of stress. For methylated guanosine derivatives (m^7^G, m^1^G, m^2^G and m^22^G), we observed a slightly reduced, but statistically significant (*e.g.* m^7^G p_6h_ = 0.0096) incorporation into tRNA^Phe^_GAA_ under stress compared to the control samples (Figure 6c and Figure S21). For Cm and Gm we observed a higher abundance under stressed conditions while m^1^A or m^5^C were comparable. Our results imply that human cells i) adapt their tRNA modifications to methylation stress by differentially modifying new transcripts and ii) consider tRNA modification as a highly important process and thus continue even during stress exposure.

## Discussion

Current analyses of the epitranscriptome are limited to snapshot moments and cannot truly follow dynamic processes inside cells. While NAIL-MS allows the observation of RNA modification adaptation processes ^37,44^ it was not possible to apply the technique in human cell culture due to the complexity of culture medium. ^13^C_6_-glucose is a reasonable and economic option for stable isotope labeling (28 € per 50 mL medium) ^45^ but it suffers from the formation of multiple isotopomers which complicates its application especially when additional feeding with CD_3_-methionine is required. In such studies, the signals of partially ^13^C-labeled nucleosides and CD_3_-methylated nucleosides can overlap and quantification becomes impossible. In contrast, supplementation of various media with _15_N_5_-adenine and ^13^C_5_15N_2_-uridine results in monoisotopic labeling with no overlap with naturally occurring ^13^C-isotopomers or artificially CD_3_-methylated nucleosides (305 € per 50 mL medium). Thus a broad applicability and even quantification by isotope dilution mass spectrometry is possible. While we observe best results with dialyzed FBS, it is also possible to use regular FBS instead if it is preferable to the cells. If the nucleoside of interest is a G or A derivative, ^13^C_6_-glucose labeling can be combined with supplementation of unlabeled adenine. This approach is less costly and produces monoisotopically labeled A and G derivatives with a ^13^C_5_-ribose moiety (Figure S3 and S4).

An important consideration for any NAIL-MS study is the constant supplementation with adenine and uridine, even when unlabeled medium is used to prevent activation of *de novo* synthesis pathways. Independently of the chosen nucleic acid labeling scheme, we strongly recommend validation experiments as shown in Figure 3c. Such an experiment is crucial to later judge the statistical significance of *e.g.* pulse chase studies. For example, our validation experiment indicates that a less than 1.6 fold change in Um would not be biologically significant (Figure S14). In such a case we recommend the direct comparison to a control sample (such as those in Figure 6) to judge the accuracy of the received NAIL-MS data.

Furthermore, we suggest careful interpretation of new transcript data at early time points of pulse chase experiments. As described for Ψ and m^5^U (Figure S15), it is possible that some modified nucleosides are early on too close to the lower limit of quantification (LLOQ) in new transcripts and thus the received quantities must be carefully interpreted.

We have studied the temporal placement of modified nucleosides in tRNA^Phe^ as a model. Our data implies that structure stabilization by modified nucleosides is a key necessity and must thus happen early on, while anticodon-loop modifications are not immediately needed and are potentially placed on-demand. One exception is the formation of wybutosine (yW). Its precursor modification m^1^G is immediately incorporated into tRNA^Phe^ before its abundance drops at later time points, presumably due to its further processing into yW. By NMR spectroscopy in combination with stable isotope labeling, Barraud *et al.* recently observed an inhibition of m^22^G formation by m^2^G ^22^. In our hands, m^22^G is placed into tRNA^Phe^ as fast as is m^2^G, but as both modifications are incorporated slowly it is possible that m^22^G is placed in a non-m^2^G modified sub-population. This question might be approached by combining NAIL with oligonucleotide MS.

With NAIL-MS we are not limited to RNA modification studies in new transcripts. In addition, we can follow the fate of RNA modifications in mature transcripts. In human cells, we observe a constant loss of modified nucleosides from tRNAs, similar to our initial report in *S. cerevisiae* ^31^. The extent of the decrease is similar for all modified nucleosides in tRNA (∼ 50 % lower within 48 h) including non-methylated modifications, which argues towards a preferential degradation of modified tRNA. In 18S rRNA, we see a similar loss of modified nucleosides from original transcripts which is with ∼ 20 % within 48 h less pronounced as in tRNA. The ∼ 2x longer half-life of rRNA compared to tRNA ^46^ supports our hypothesis of preferred degradation of modified RNA which is most likely connected to the life time of RNA.

The constant loss of pre-existing modifications from original RNA is masked in the early time points of the experiment by observable post-transcriptional methylation of original RNA. For many modified nucleosides, the extent of post-methylation of existing transcripts is connected to the extent of modification in new transcripts (Figure S17). Some modified nucleosides such as m^7^G, m^3^U, m^3^C, mcm^5^s^2^U and Um show no correlation between post-methylation and new methylation abundance. Except m^7^G, all these modified nucleosides are placed in or close to the anticodon-loop which indicates that the modification extent at these positions reflects rather demand than maturation. Another hypothesis for the post-methylation arises from reports on tRNA demethylation. For m^1^A and m^3^C, demethylation by members of the ALKBH family has been proposed ^9,47,48^. Such a demethylated site might be target to re-methylation and this process would lead to the formation of post-methylated nucleosides. While the common analysis of tRNA modifications by sequencing and quantitative mass spectrometry provides a static view on the substrates of ALKBH enzymes, future NAIL-MS experiments will shed light onto the dynamic performance of these enzymes *in vivo*.

Such a detailed analysis is especially important for understanding the processes behind stress induced adaptation of tRNA modifications. To this end, we have studied the impact of methylation stress on tRNA modifications. Even at a harsh dose of MMS (1 mM), we observe only 1 damage derived m^1^A and 5 m^7^G per 1000 tRNAs. Other damage products were not observed. Intriguingly, these damages do not seem to be repaired in human cells.

In our hands, methylation stress has no impact on the abundance of modified nucleosides in tRNA present during the stress exposure. In contrast, the abundance of some modified nucleosides is slightly, but significantly changed in new transcripts. This indicates that cells regulate their tRNA modifications on the level of new transcripts and not existing transcripts. Overall modification processes of tRNA are not stalled during stress recovery which indicates that properly modified tRNAs are of high importance to the cell.

NAIL-MS is a powerful technique which depends, as common to state-of-the-art mass spectrometry of modified nucleosides, on a complete enzymatic digest to the nucleoside building block. Thus all sequence context surrounding modified nucleosides is lost and the technique relies strongly on the purity of the sample. This is especially important for mRNA ^49^. If reliable mRNA purification is possible, the true dynamics of m^6^A and other mRNA modifications becomes finally available through NAIL-MS.

## Acknowledgement

This study was funded through the Deutsche Forschungsgemeinschaft (KE1943/3-1). M.H., F.H. and S.K. thank Angie Kirchner, Thomas Carell and his group for instrument time (high-resolution mass spectrometer) and advice.

## Author contribution

M.H. and S.K. planned the experiments and wrote the manuscript. M.H. and F.H. conducted the experiments. M.H. and S.K. performed data analysis.

## Additional information

Supplementary information is available. Correspondence and requests for materials should be addressed to S.K..

## Data availability

The data that supports the findings of this study are available from the corresponding author upon reasonable request.

## MATERIAL & METHODS

### Salts, reagents, media and nucleosides

All salts, reagents and media were obtained from Sigma-Aldrich (Munich, Germany) at molecular biology grade unless stated otherwise. The isotopically labeled compounds ^13^C_5_,^15^N_2_-Uridine (Ribose-^13^C_5_, 98%; ^15^N_2_, 96-98%) and ^15^N_5_-Adenine (^15^N_5_, 98%) were obtained from Cambridge Isotope Laboratories (Tewksbury, MA, USA). Unlabeled glutamine, isotopically labeled L-glutamine-amide-^15^N (98 atom% ^15^N), L-aspartic-^15^N acid (98 atom% ^15^N) and (D_3_)-L-methionine (98 atom% D) were obtained from Sigma-Aldrich. Isotopically labeled ^13^C_6_-glucose (≥99 atom% ^13^C) was obtained from Eurisotope (Saarbruecken, Germany). All solutions and buffers were made with water from a Sartorious arium^®^ pro ultrapure water system (Goettingen, Germany). The nucleosides adenosine (A), cytidine (C), guanosine (G) and uridine (U), were obtained from Sigma-Aldrich. 1-Methyladenosine (m^1^A), N3-methylcytidine (m^3^C), N6-methyladenosine (m^6^A), 7-methylguanosine (m^7^G), 5-methylcytidine (m^5^C), 5-methyluridine (m^5^U), 2’-O-methylcytidine (Cm), 2’-O-methylguanosine (Gm), 1-methylguanosine (m^1^G), N2-methylguanosine (m^2^G), 2-dimethylguanosine (m^22^G), pseudouridine (Ψ), inosine (I), 2’-O-methyluridine (Um), 2’-O-methyladenosine (Am) and 5-methoxycarbonylmethyl-2-thiouridine (mcm^5^s^2^U) were obtained from Carbosynth (Newbury, UK). Dihydrouridine (D) was obtained from Apollo Scientific (Stockport, UK). N6-threonylcarbamoyladenosine (t^6^A) was obtained from TRC (North York, Canada). N3-methyluridine (m^3^U) and N6-isopentenyladenosine (i^6^A) were generous gifts from the Dedon lab. 5-carbamoylmethyl-2-thiouridine (ncm^5^s^2^U) was a generous gift from the Helm lab. 1-Methylinosine (m^1^I) was a generous gift from STORM Therapeutics LTD (Cambridge, UK).

### Cell culture

All cell culture media and supplements were obtained from Sigma-Aldrich (Munich, Germany) unless stated otherwise. Standard Basal medium for HEK 293 culture was DMEM D6546 high glucose supplemented with 10% FBS and 0.584 g/L L-glutamine. Cells were split 1:7 using standard procedures every 2-3 days to counter overgrowth. Cells cultured in DMEM medium were kept at 10% CO_2_ for proper pH adjustment. For all experiments where labeling of nucleosides was involved DMEM D0422 without methionine and cysteine was used. DMEM D0422 was supplemented with 10% dialyzed FBS (Biowest, Nuaillé, France), 0.584 g/L L-glutamine, 0.063 g/L cystine (stock concentration 78.75 g/L dissolved in 1M HCl), 0.03 g/L methionine, 0.05 g/L uridine and 0.015 g/L adenine. Uridine, adenine and methionine were either added as unlabeled or labeled compounds depending on the desired labeling. HeLa cells were cultured and labeled using the same media.

For Labeling in RPMI R0883, dialyzed FBS, glutamine, methionine, uridine and adenine were added in the same concentrations as for DMEM D0422.

HAP1 cells were either labeled using DMEM D0422 as described above or IMDM I3390 where FBS, glutamine, uridine, adenine and methionine were added in the same concentrations as used for DMEM D0422 medium. Cells grown in RPMI or IMDM medium were kept at 5% CO_2_ for proper pH adjustment.

Mouse embryonic stem cells (mESC) were cultured as recently reported ^50^. Isotopically labeled compounds were added as described for regular cell culture labeling.

For biological replicates one culture was split into several flask at least 24h prior to experiment initiation.

### Cell lysis and RNA purification

Cells were directly harvested on cell culture dishes using 1 mL TRI reagent for T25 flasks or 0.5 mL TRI reagent for smaller dishes. The total RNA was isolated according to the supplier’s manual with chloroform (Roth, Karlsruhe, Germany). tRNA and 18S rRNA were purified by size exclusion chromatography (AdvanceBio SEC 300Å, 2.7μm, 7.8×300mm for tRNA and BioSEC 1000Å, 2.7μm, 7.8×300mm for 18S rRNA, Agilent Technologies) according to published procedures ^31,51^. The RNA was resuspended in water (35 μL).

### Isoacceptor purification

The procedure was adapted from Hauenschild *et al*. ^13^. For tRNA^Phe^_GAA_ purification, 1 μg pre-purified total tRNA was used. The sequence of the biotinylated 2’-deoxyoligonucleotide is 5’ – (Biotin) AAATGGTGCCGAAACCCGGGATCGAACCAGGGT – 3’ (Sigma Aldrich, Munich, Germany)

### tRNA digestion for mass spectrometry

Total tRNA (300 ng) in aqueous digestion mix (30 μL) was digested to single nucleosides by using 2 U alkaline phosphatase, 0.2 U phosphodiesterase I (VWR, Radnor, Pennsylvania, USA), and 2 U benzonase in Tris (pH 8, 5 mM) and MgCl_2_ (1 mM) containing buffer. Furthermore, 0.5 µg tetrahydrouridine (Merck, Darmstadt, Germany), 1 µM butylated hydroxytoluene, and 0.1 µg pentostatin were added to avoid deamination and oxidation of the nucleosides. When quantification of dihydrouridine was intended tetrahydrouridine was omitted. After incubation for 2 h at 37 °C, 20 µL of LC-MS buffer A (QQQ) was added to the mixture and then filtered through 96-well filter plates (AcroPrep Advance 350 10 K Omega, PALL Corporation, New York, USA) at 3000 ×g and 4 °C for 30 min. A stable isotope labeled internal standard (SILIS) was produced in *S. cerevisiae* using ^13^C and ^15^N rich growth medium (Silantes, Munich, Germany, Product# 111601402) following recently described procedures ^16,31^. 1/10 Vol. of SILIS was added to each filtrate before analysis by QQQ mass spectrometry. For each sample 10 µL were injected (∼90 ng of sample tRNA)

### High resolution mass spectrometry

The ribonucleosides were separated using a Dionex Ultimate 3000 HPLC system with a Synergi, 2.5 μm Fusion-RP, 100 Å, 100 x 2 mm column (Phenomenex®, Torrance, California, USA). Mobile phase A was 10 mM ammonium formate and mobile phase B was 80% acetonitrile containing 2 mM ammonium formate. Gradient elution started with 0 % B and increased to 12% B after 10 min and to 80% after 12 min. After 4 min elution at 80 % B and subsequently regeneration of starting conditions to 100% A after 5 min, the column was equilibrated at 100% A for 8 min. The flow rate was 0.2 mL/min and the column temperature 30 °C. High-resolution mass spectra were recorded by a ThermoFinnigan LTQ Orbitrap XL operated in positive ionization mode. The parameters of the mass spectrometer were tuned with a freshly mixed solution of uridine (10 μM). Capillary voltage was set to 20 V and capillary temperature to 300 °C. Sheath gas and sweep gas flow rate was set to 0, and auxiliary gas flow rate to 35. Source voltage was set to 4.0 kV and tube lens to 75 V.

### QQQ mass spectrometry

For quantitative mass spectrometry an Agilent 1290 Infinity II equipped with a diode-array detector (DAD) combined with an Agilent Technologies G6470A Triple Quad system and electrospray ionization (ESI-MS, Agilent Jetstream) was used. Operating parameters: positive-ion mode, skimmer voltage of 15 V, cell accelerator voltage of 5 V, N_2_ gas temperature of 230 °C and N_2_ gas flow of 6 L/min, sheath gas (N_2_) temperature of 400 °C with a flow of 12 L/min, capillary voltage of 2500 V, nozzle voltage of 0 V, and nebulizer at 40 psi. The instrument was operated in dynamic MRM mode (Table S2).

For separation a Synergi, 2.5 μm Fusion-RP, 100 Å, 100 x 2 mm column (Phenomenex®, Torrance, California, USA) at 35 °C and a flow rate of 0.35 mL/min was used in combination with a binary mobile phase of 5 mM NH_4_OAc aqueous buffer A, brought to pH 5.6 with glacial acetic acid (65 μL/L), and an organic buffer B of pure acetonitrile (Roth, Ultra LC-MS grade, purity ≥99.98). The gradient started at 100% solvent A for 1 min, followed by an increase to 10% solvent B over 4 min. From 5 to 7 min, solvent B was increased to 40% and maintained for 1 min before returning to 100 % solvent A in 0.5 min and a 2.5 min re-equilibration period.

### Calibration

For calibration, synthetic nucleosides were weighed and dissolved in water to a stock concentration of 1-10 mM. The calibration solutions ranged from 0.025 to 100 pmol for each canonical nucleoside and from 0.00125 pmol to 5 pmol for each modified nucleoside. Each calibration was spiked with 10% SILIS. The sample data were analyzed by the quantitative and qualitative MassHunter Software from Agilent. The areas of the MRM signals were integrated for each modification. The values of integrated MS signals from target nucleosides were set into relation to the respective MS signals of the respective isotope labeled SILIS nucleosides after Equation **(1)** to receive the nucleoside isotope factor (NIF):

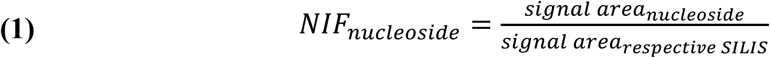

Results from Equation 1 were plotted against the expected molar amount of nucleosides and regression curves were plotted through the data points. The slopes represent the respective relative response factors for the nucleosides (rRFN) and enable an absolute quantification. The principle is described in more detail in our published protocol ^16^. The plotting of these calibration curves is done automatically by the quantitative MassHunter Software and should be checked manually for linearity.

### Data Analysis

Molar amounts of nucleosides in samples were calculated after Equation **(2)** using the signal areas of target compounds and SILIS in the samples and the respective rRFN, determined by calibration measurements. This step is done automatically by the quantitative MassHunter Software.

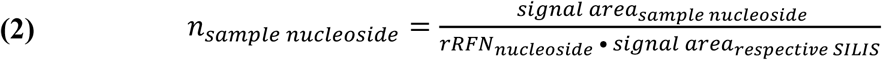

To make different samples quantitatively comparable, the molar amount of each modified nucleoside was normalized by the molar amount of injected RNA to receive the number of modifications per RNA. Therefore, the calculated amounts of injected canonicals were divided by their expected occurrence in the respective RNAs and averaged afterwards (see Equation **(3)** for tRNA). The numbers for each canonical nucleoside were either taken from the sequence of 18S rRNA, tRNA^Phe^ (reported modifications subtracted)^32^ or determined empirically for total tRNA analyses.

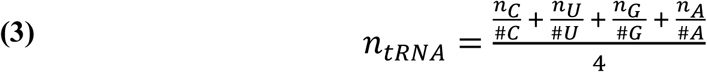

In the case of NAIL-MS experiments, the different isotopomers were referenced to their respective labeled canonicals, so that original (unlabeled) modifications were referenced to original tRNA molecules and new (labeled) modifications were referenced to new tRNA molecules (see Equations **(4)** and **(5)**). Table 1 gives a summary of the calculations exemplarily for m^7^G.

**Table 1:** Quantification of m^7^G per tRNA (based on G) First the molar amount of injected nucleosides is calculated based on the signal areas of target nucleosides and SILIS and the respective rRFNs determined by calibration (here for m^7^G and G). Then the molar amount of modification is divided by the molar amount of respective tRNA calculated by dividing the molar amount of canonical by the expected number of the respective canonical (here just based on G).

|  | m<sup>7</sup>G (pmol) | G (pmol) | m<sup>7</sup>G per tRNA |
| --- | --- | --- | --- |
| **original** | $\frac{\text{area m}^7\text{G (unlabeled)}}{\text{rRFN m}^7\text{G} \cdot \text{area m}^7\text{G (SILIS)}}$ | $\frac{\text{area G (unlabeled)}}{\text{rRFN G} \cdot \text{area G (SILIS)}}$ | $\frac{\text{m}^7\text{G (original)}}{\frac{\text{G (original)}}{\# \text{ of G in sequence}}}$ |
| **new** | $\frac{\text{area m}^7\text{G (}^{15}\text{N,CD}_3\text{)}}{\text{rRFN m}^7\text{G} \cdot \text{area m}^7\text{G (SILIS)}}$ | $\frac{\text{area G (}^{15}\text{N)}}{\text{rRFN G} \cdot \text{area G (SILIS)}}$ | $\frac{\text{m}^7\text{G (new)}}{\frac{\text{G (new)}}{\# \text{ of G in sequence}}}$ |

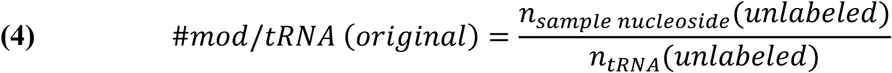

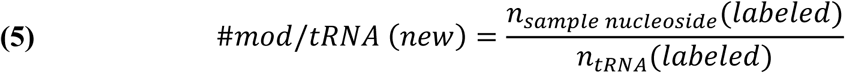

### Statistics

All experiments were performed at least 3 times (biological replicates) to allow student t-test analysis. P-values of student t-test (unpaired, two-tailed, equal distribution) were calculated using Excel.

