## Supplement Information for "Cell culture NAIL-MS allows insight into human RNA modification dynamics *in vivo*"

Matthias Heiss[1], Felix Hagelskamp[1] and Stefanie Kellner[1]*
[1] Department of Chemistry, Ludwig-Maximilians-University Munich, Butenandtstr. 5-13, 81377 Munich, Germany


* To whom correspondence should be addressed

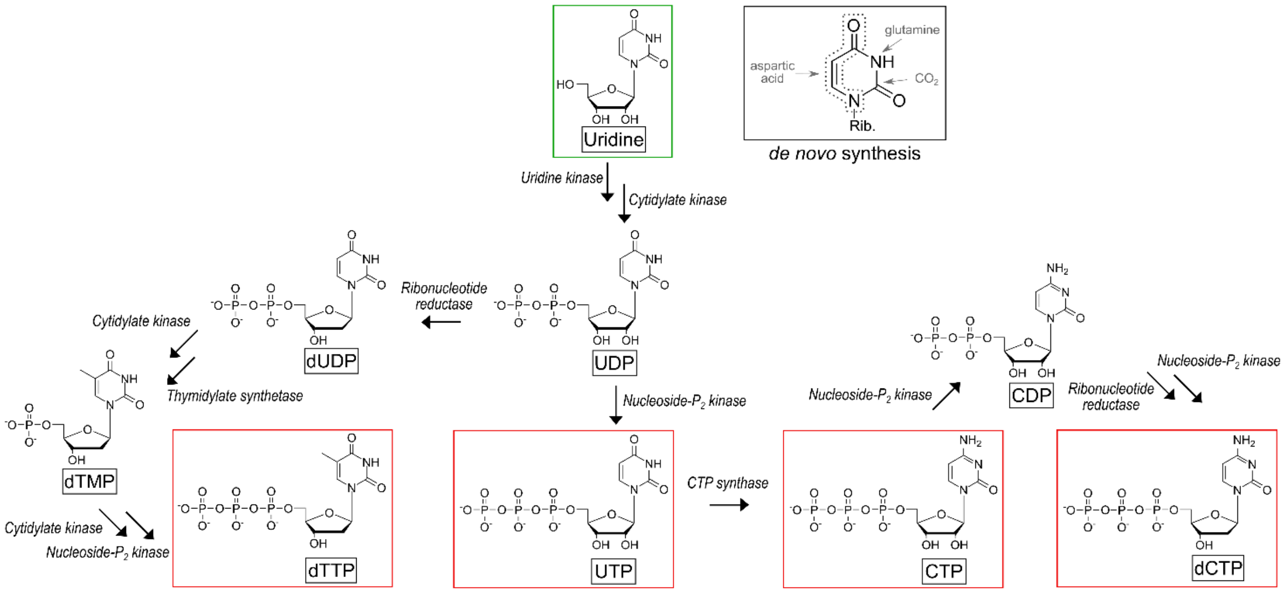


**Figure S1A: Biosynthesis and connection of pyrimidines**

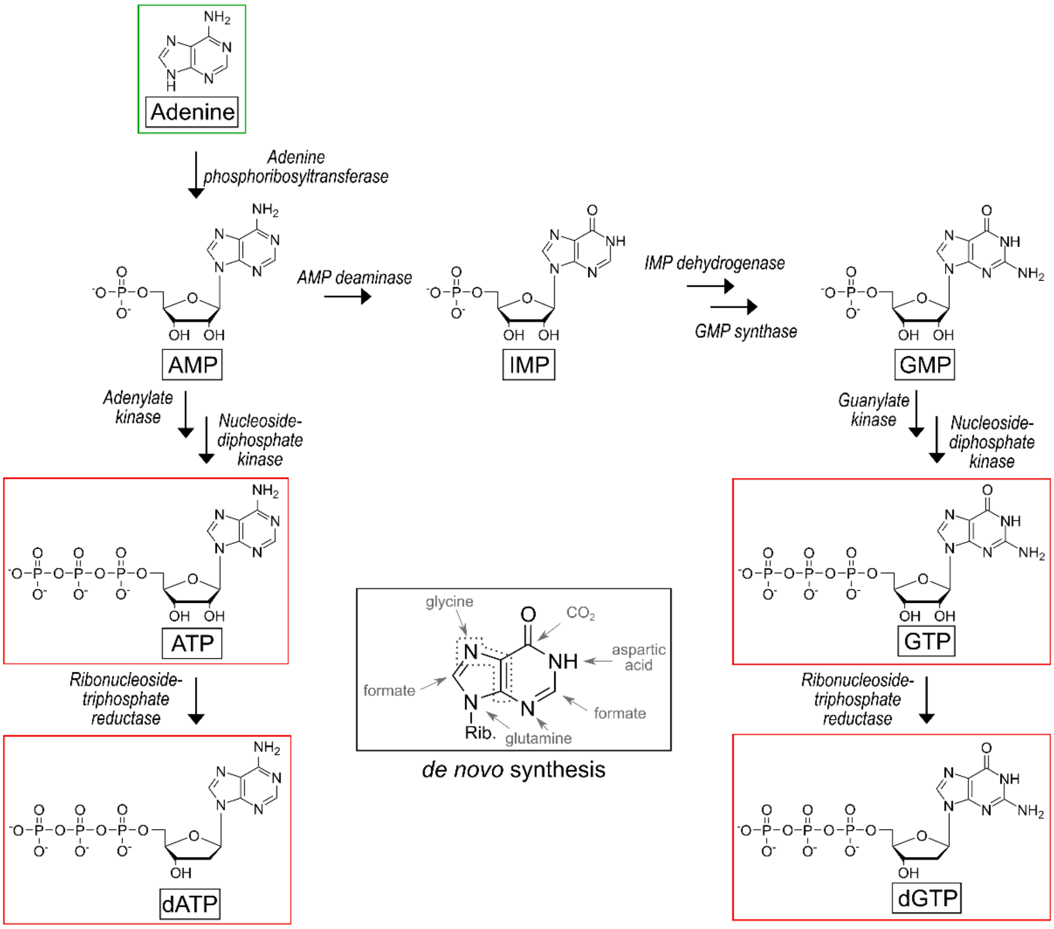


**Figure S1B: Biosynthesis and connection of purines**

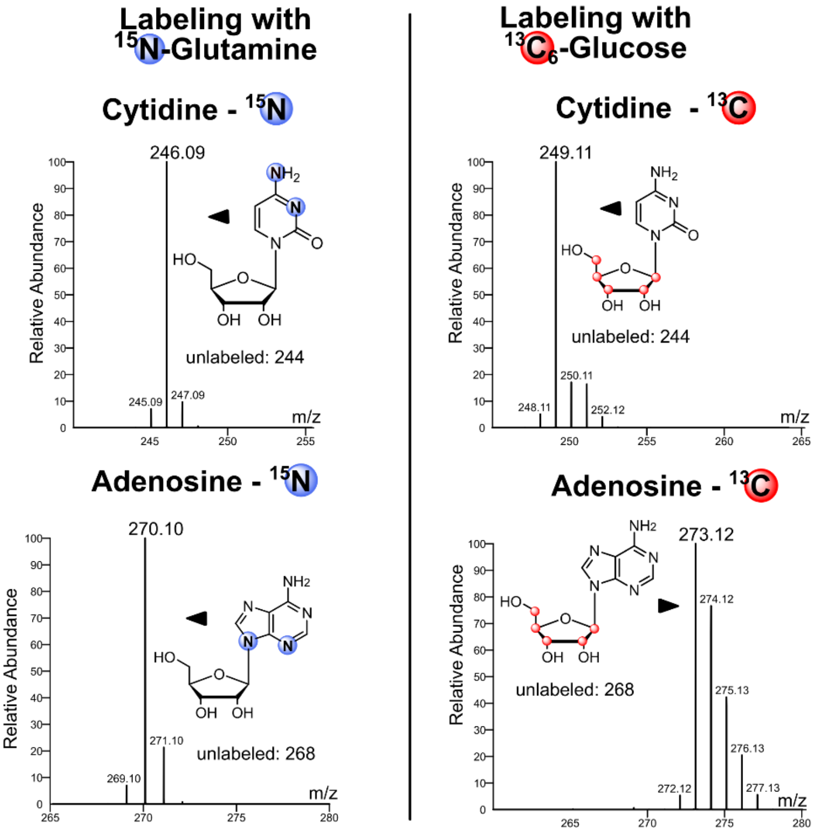


**Figure S1C: Labeling of nucleosides with $^{15}N_2$-glutamine and $^{13}C_6$-glucose.**
The metabolites used for biosynthesis of pyrimidines and purines are shown on the left. The high-resolution mass spectra of cytidine (top) and adenosine (bottom) from isolated RNA after labeling with $^{15}N_2$-glutamine or $^{13}C_6$-glucose are shown on the right. All cells were labeled for 7 days (=3 passages) in the respective medium.

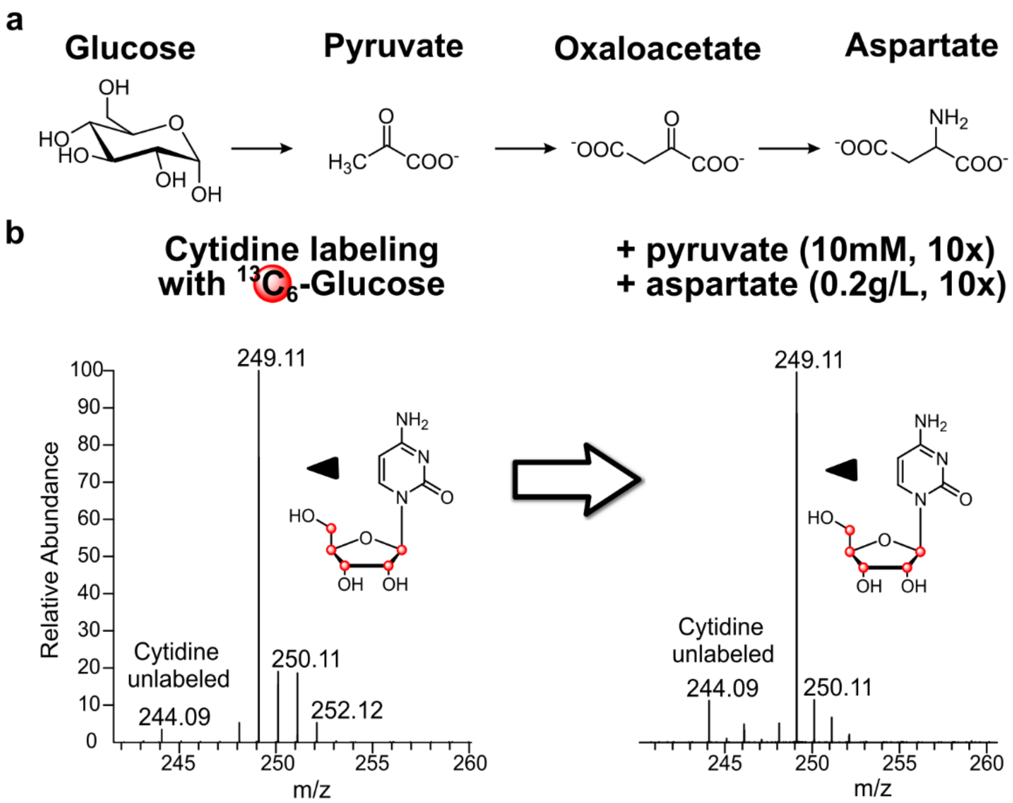


**Figure S2: Pyrimidine labeling in $^{13}C_6$-glucose medium supplemented with pyruvate and aspartate.**

**a,** Reaction scheme showing the biosynthesis pathway of aspartate with pyruvate as an intermediate. **b,** Left: High-resolution mass spectrum of cytidine after labeling with $^{13}C_6$-glucose. Right: High-resolution mass spectrum of cytidine with additional supplementation of pyruvate and aspartate to overcome poly-isotopic labeling. All cells were labeled for 7 days (=3 passages) in the respective medium.

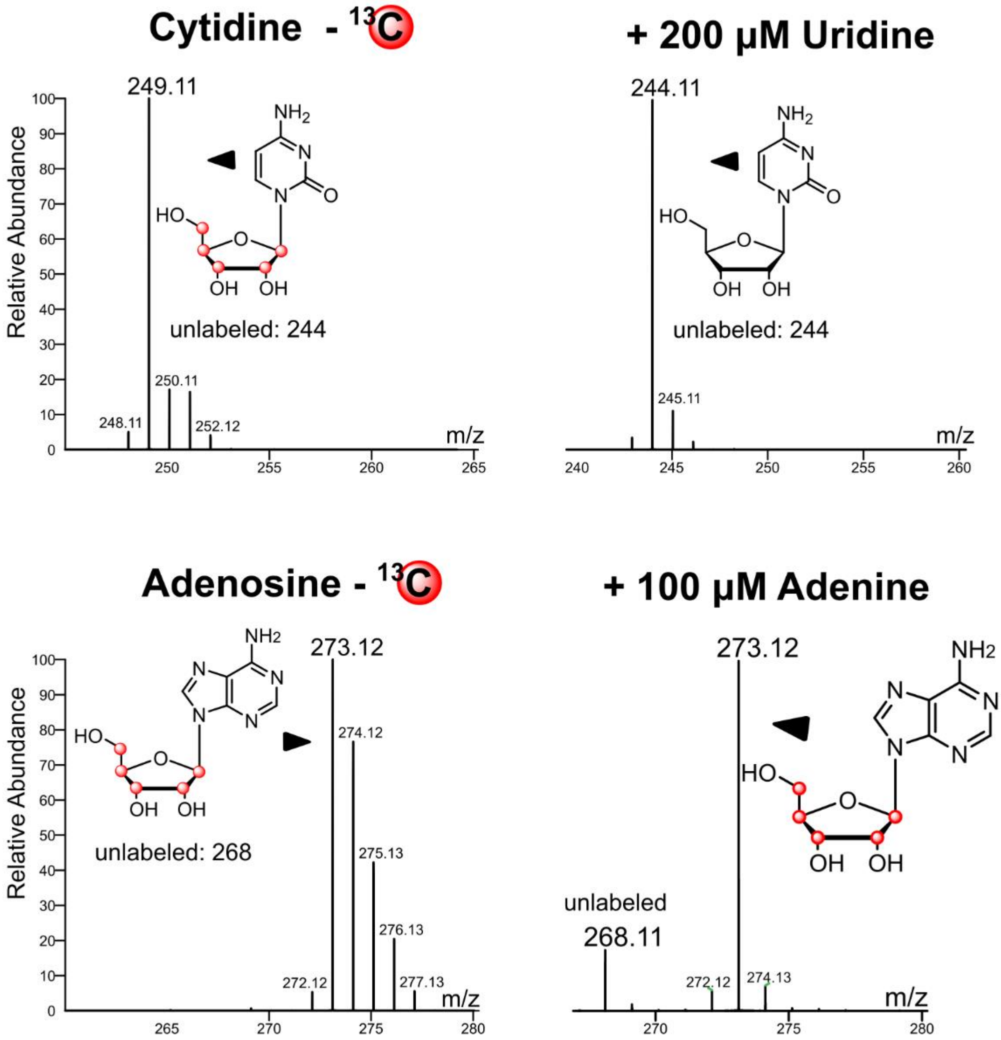


**Figure S3: Improvement of nucleoside labeling in $^{13}C_6$-glucose medium by supplementation of uridine and adenine.**

The high-resolution mass spectra on the left show labeling of cytidine (top) and adenosine (bottom) after labeling with $^{13}C_6$-glucose. The spectra on the right show the mono-isotopic signals after additional supplementation with 200 µM uridine and 100 µM adenine. All cells were labeled for 7 days (=3 passages) in the respective medium.

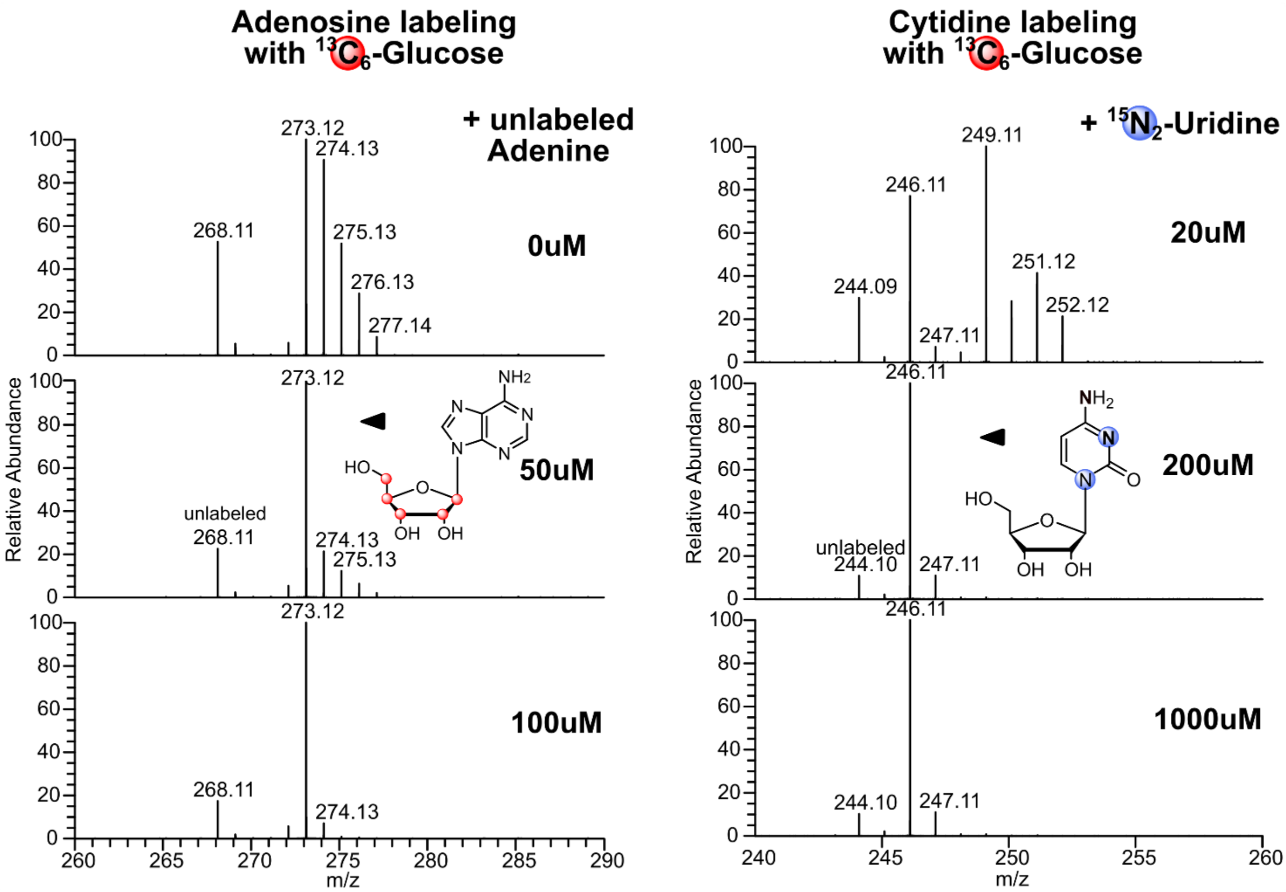


**Figure S4: Concentration optimization of adenine and uridine for stable isotope labeling.**
The high-resolution mass spectra on the left show stable isotope labeling of adenosine after labeling with $^{13}C_6$-glucose and different concentrations of unlabeled adenine. The high-resolution spectra on the right show stable isotope labeling of cytidine after labeling with $^{13}C_6$-glucose and different concentrations of $^{15}N_2$-uridine. All cells were labeled for 7 days (=3 passages) in the respective medium. Note: Unlabeled signals are caused by the low labeling efficiency in $^{13}C_6$-glucose medium.

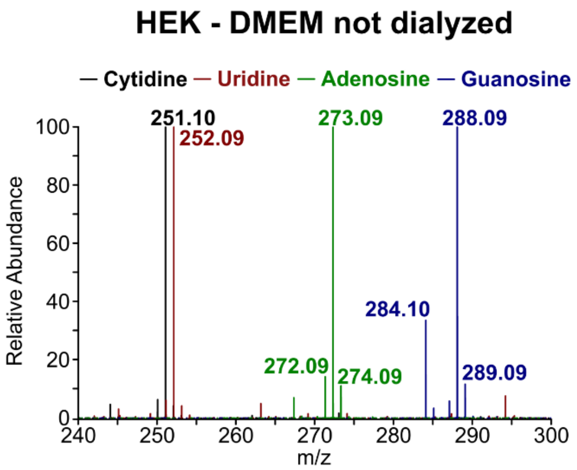


**Figure S5: Stable isotope labeling of nucleosides with labeled uridine and adenine using undialyzed FBS.**
Merged high-resolution mass spectra of cytidine, uridine, guanosine and adenosine after stable isotope labeling of HEK 293 cell culture using DMEM D0422 supplemented with $^{13}C_5$,$^{15}N_2$-uridine and $^{15}N_5$-adenine but normal FBS instead of dialyzed FBS. Cells were labeled for 7 days (=3 passages).

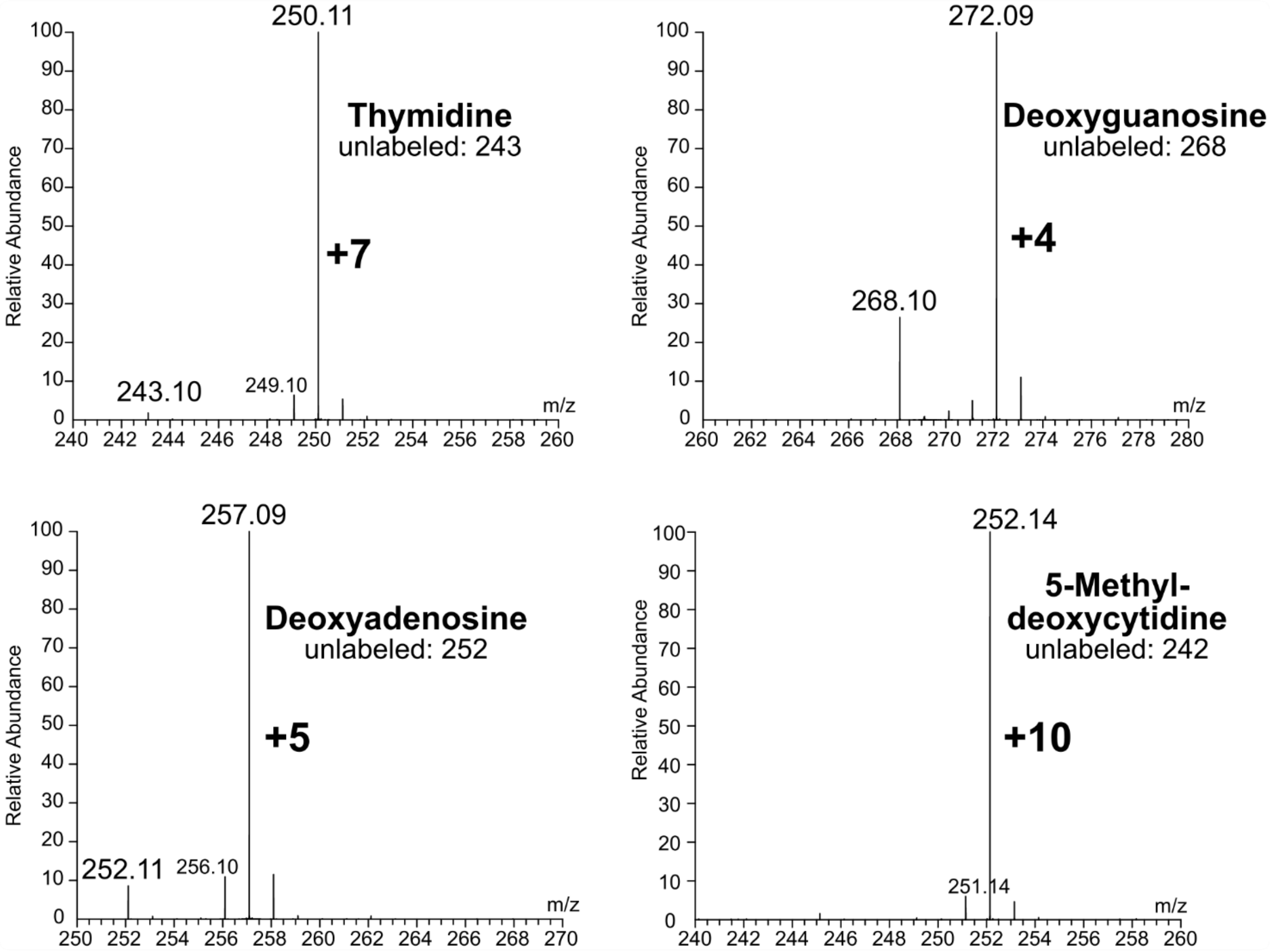


**Figure S6: Stable isotope labeling of DNA**

High-resolution mass spectra of DNA nucleosides after labeling in DMEM D0422 medium supplemented with stable isotope labeled $^{15}N_5$-adenine, $^{13}C_5{}^{15}N_2$-uridine and $CD_3$-methionine. Cells were labeled for 7 days (=3 passages) and DNA was purified (mini spin columns) and digested to nucleosides using standard procedures.

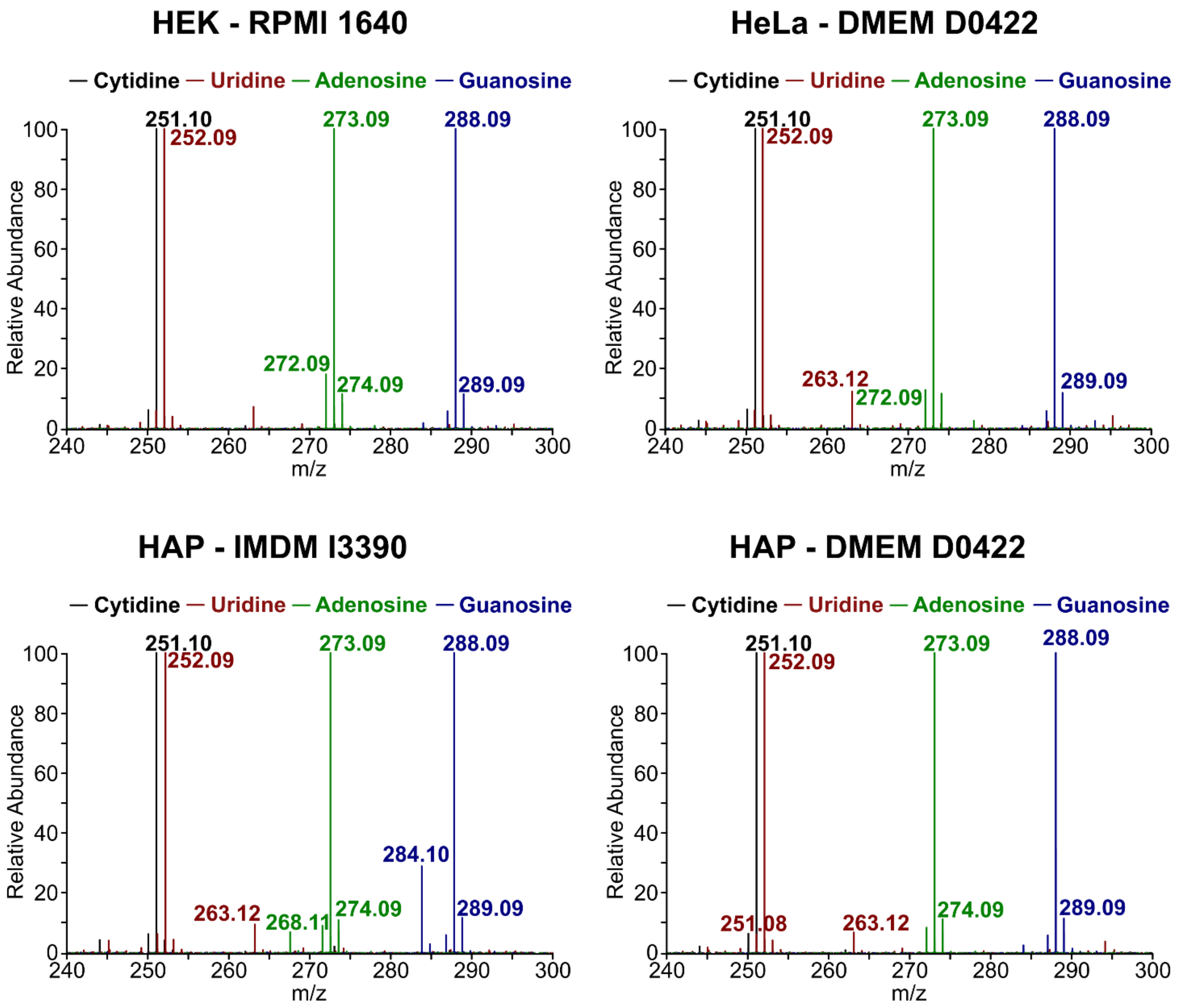


**Figure S7: Stable isotope labeling of different cell lines in different media.**
Merged high-resolution spectra of cytidine, uridine, guanosine and adenosine after stable isotope labeling of different cell lines using the respective medium supplemented with stable isotope labeled $^{15}N_5$-adenine and $^{13}C_5$,$^{15}N_2$-uridine and dialyzed FBS. All cells were labeled for 7 days (=3 passages) in the respective medium.

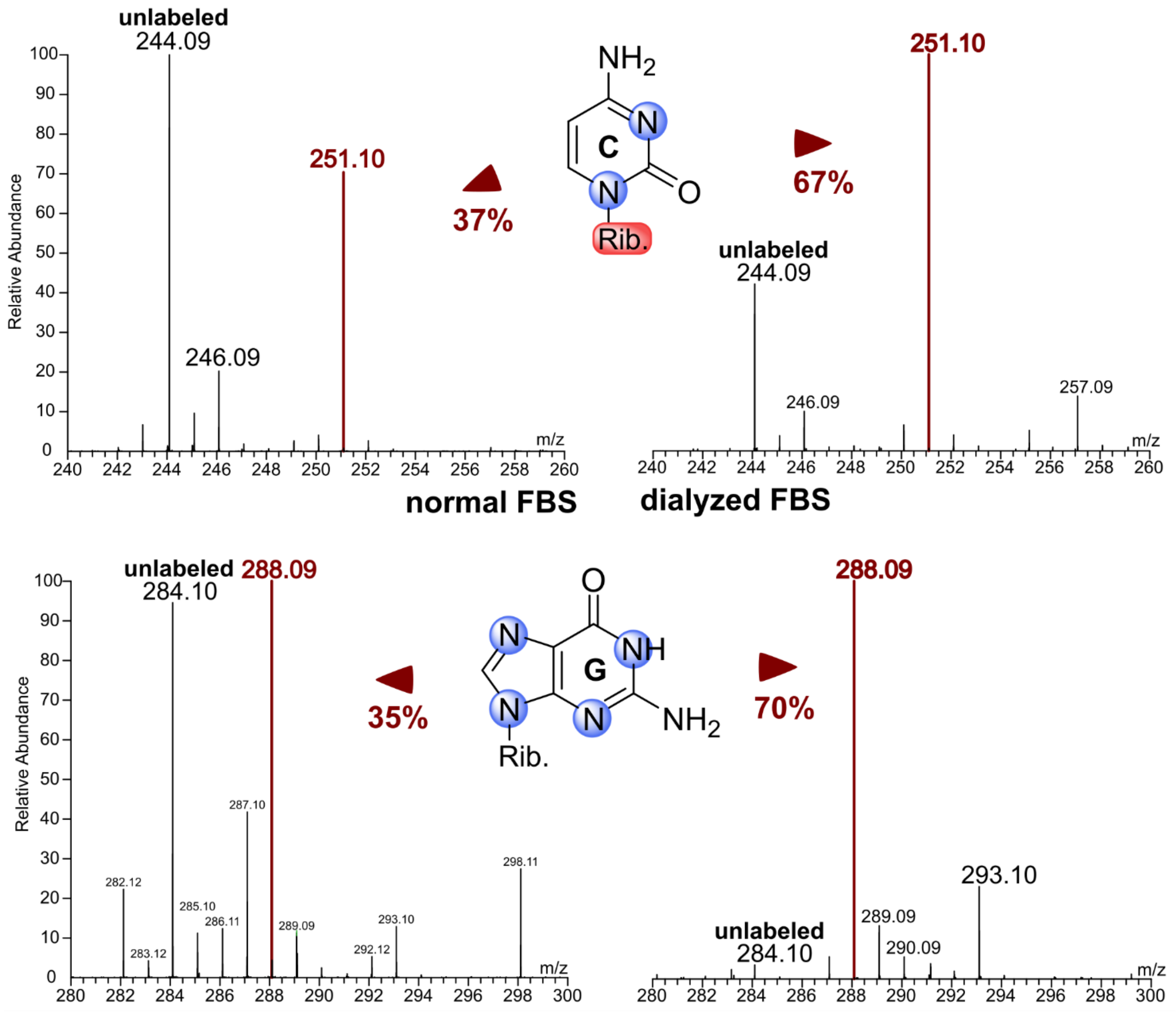


**Figure S8: Labeling of mESC RNA.**

The high-resolution mass spectra on the left show stable isotope labeling of cytidine (top) and guanosine (bottom) after labeling mouse embryonic stem cells with $^{15}N_5$-adenine and $^{13}C_5{}^{15}N_2$-uridine in DMEM containing FBS and LIF (leukemia inhibitory factor) for 4 days (for detailed information see Rahimoff *et al.* [1]. The percentages were calculated by adding up the relative abundances of all peaks for the nucleoside of interest. The relative abundance of the peak with desired labeling was then divided by the sum of all peaks.

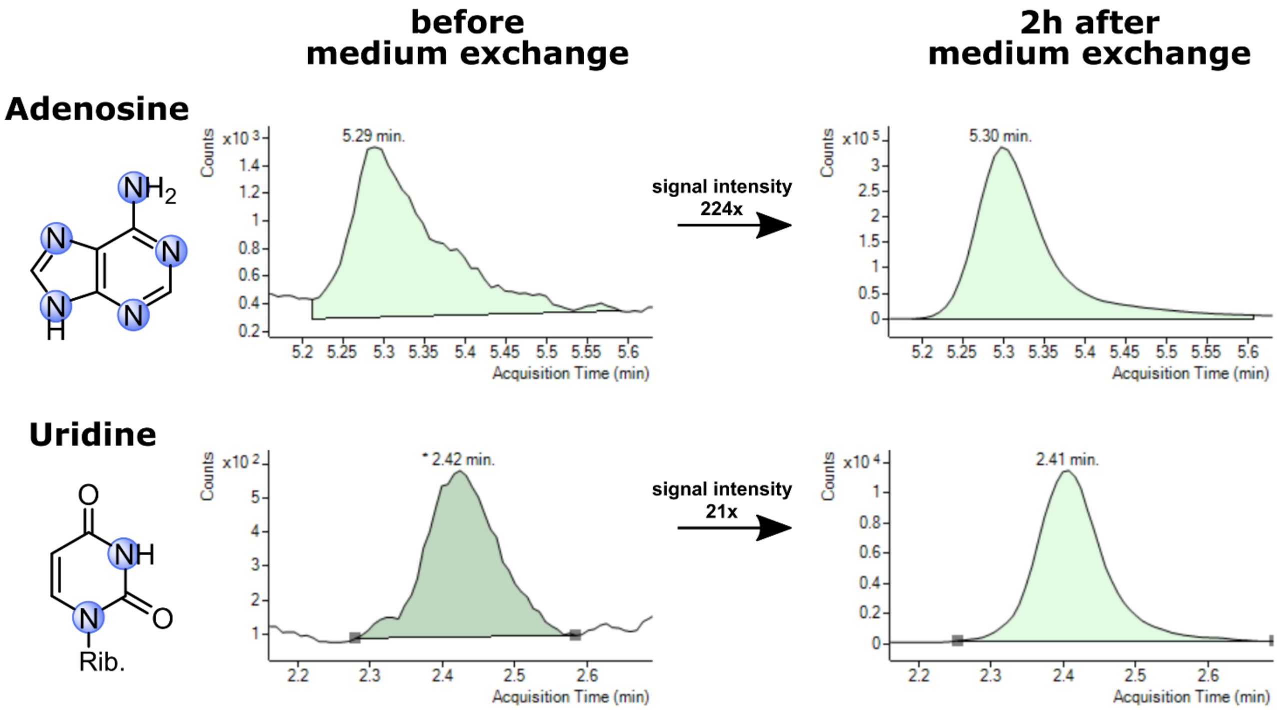


**Figure S9: Occurrence of labeled nucleosides 2 hours after growth to labeled media.**
MS signals of labeled adenosine (top, 273 → 136) and uridine (bottom, 247 →115) without stable isotope labeling are shown on the left. MS signals of labeled adenosine and uridine after labeling for 2 hours are shown on the right.

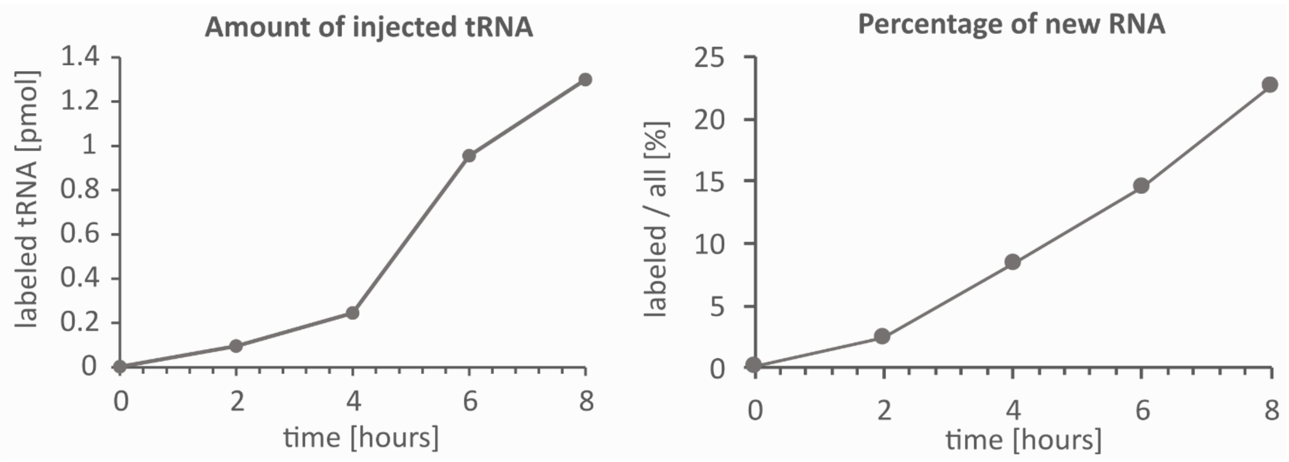


**Figure S10: Increase of labeled tRNA molecules.**
For all time points the same mass of RNA was injected (according to UV measurements of undigested RNA using an IMPLEN Nanophotometer, Munich, Germany). The injected amount of tRNA was calculated based on the sum of measured values for C, U, G and A. Left: Absolute amount of labeled tRNA. Right: Relative increase of labeled tRNA in proportion to all tRNA (unlabeled + labeled).

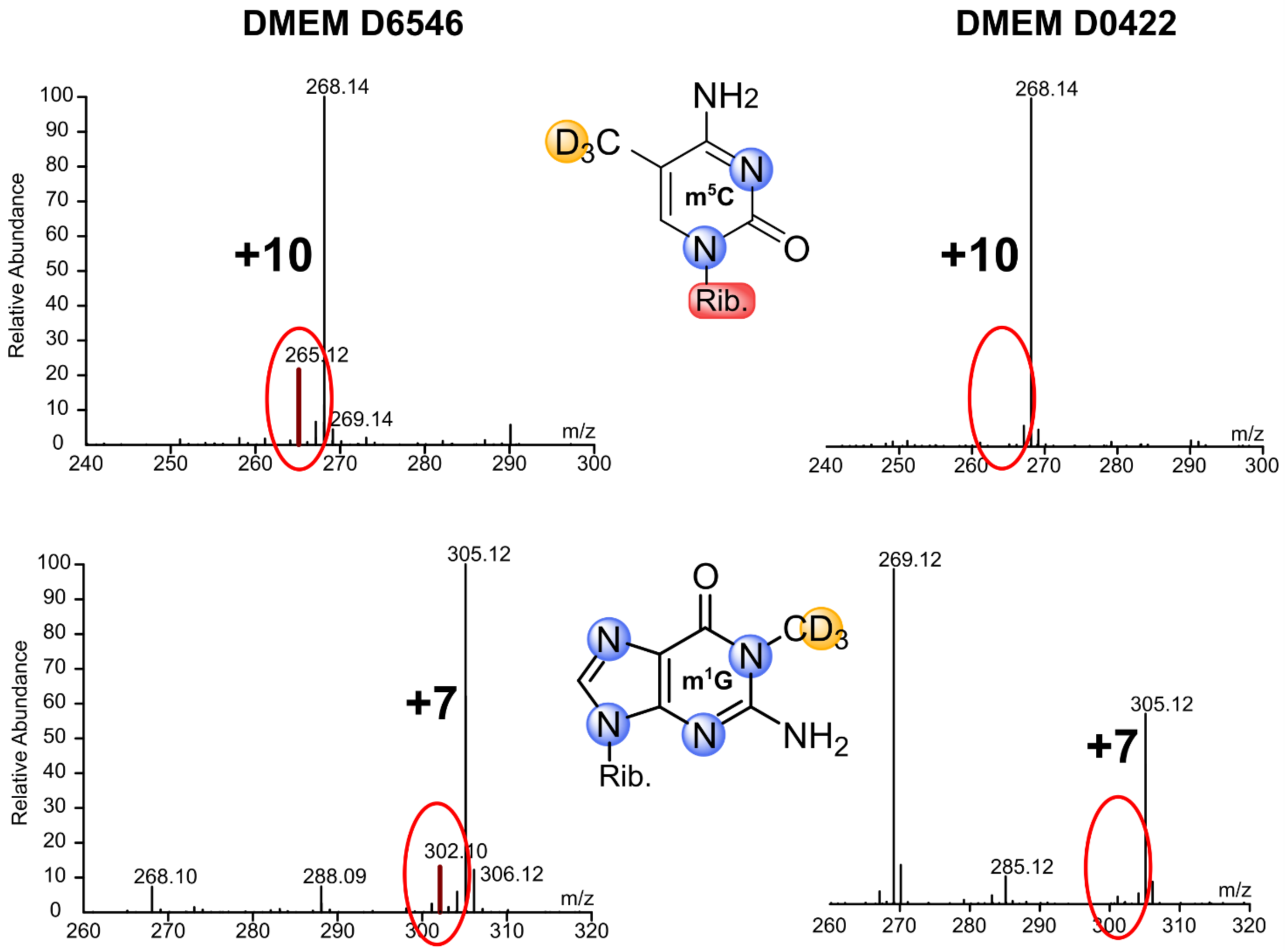


**Figure S11: Labeling of methyl-groups by $D_3$-methionine.**
Left: High-resolution mass spectra of $m^5C$ and $m^1G$ after labeling in DMEM D6546 supplemented with stable isotope labeled $^{15}N_5$-adenine and $^{13}C_5{}^{15}N_2$-uridine and 0.15 g/L (5x) $D_3$-methionine. Right: High-resolution mass spectra of $m^5C$ and $m^1G$ after labeling in DMEM D0422 supplemented with stable isotope labeled $^{15}N_5$-adenine and $^{13}C_5{}^{15}N_2$-uridine and 0.03 g/L (1x) $D_3$-methionine. All cells were labeled for 7 days (=3 passages) in the respective medium. Red circles highlight the signal for $^{13}C/^{15}N$-labeled nucleosides with an undesired $CH_3$-methylation.

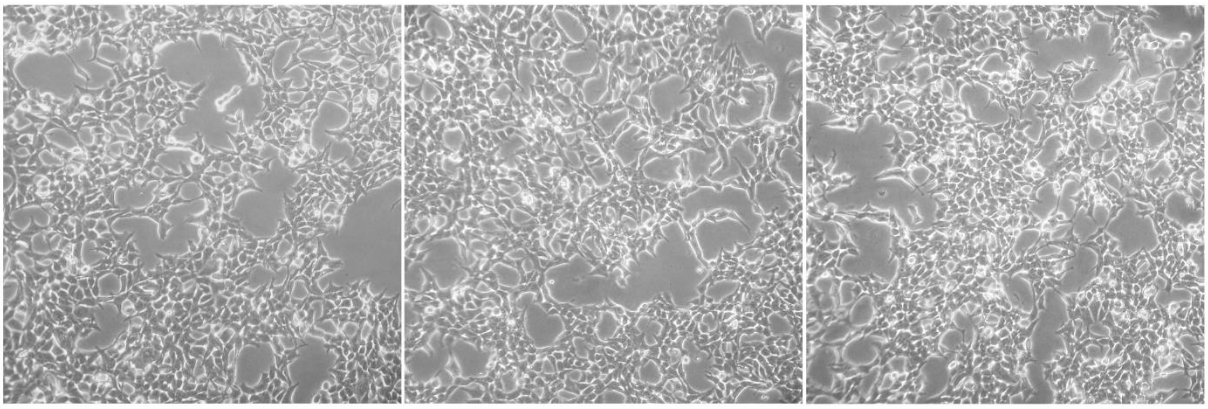


**Figure S12: Photographs of HEK 293 cells grown in different DMEM media.**
1.2 million cells were plated in a T25 flask and cultured in the respective media for 2 days. DMEM D6546 was supplemented with glutamine and FBS only. DMEM D0422 was supplemented with glutamine, FBS and cystine. Methionine, adenine and uridine were supplemented in DMEM D0422 as unlabeled or labeled compounds respectively.

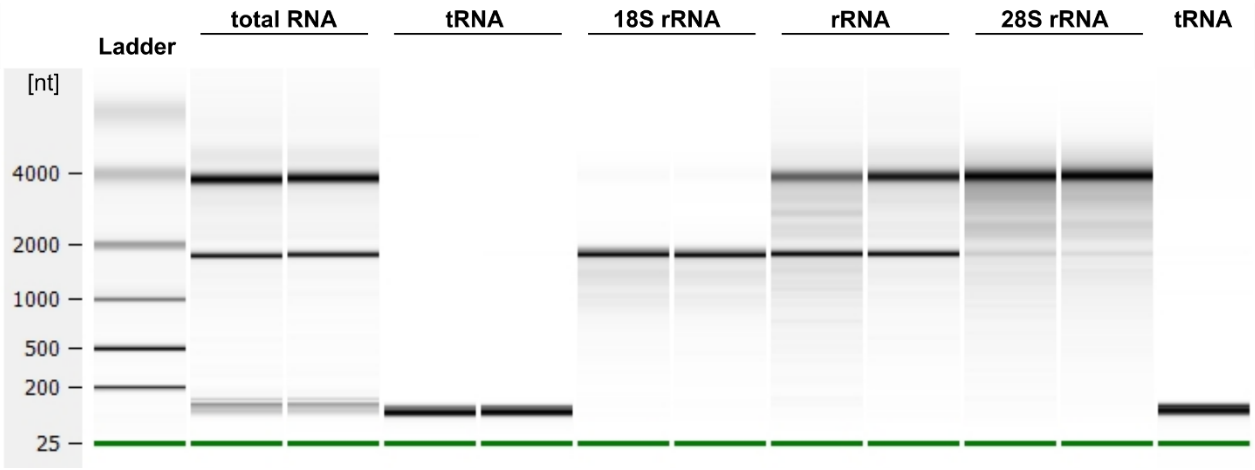


**Figure S13: BioAnalyzer Pico chip of SEC purified RNA species.** For each species the exemplary timepoint = 0 of a forward and a reverse experiment are shown. Total RNA, 18S rRNA, total rRNA and 28S rRNA are from samples shown in Fig. 3. tRNA is from samples shown in Fig. 4 and was used for purification of $tRNA^{Phe}$. tRNA in the right lane was used for purification of $tRNA^{Phe}$ for Fig. 1.

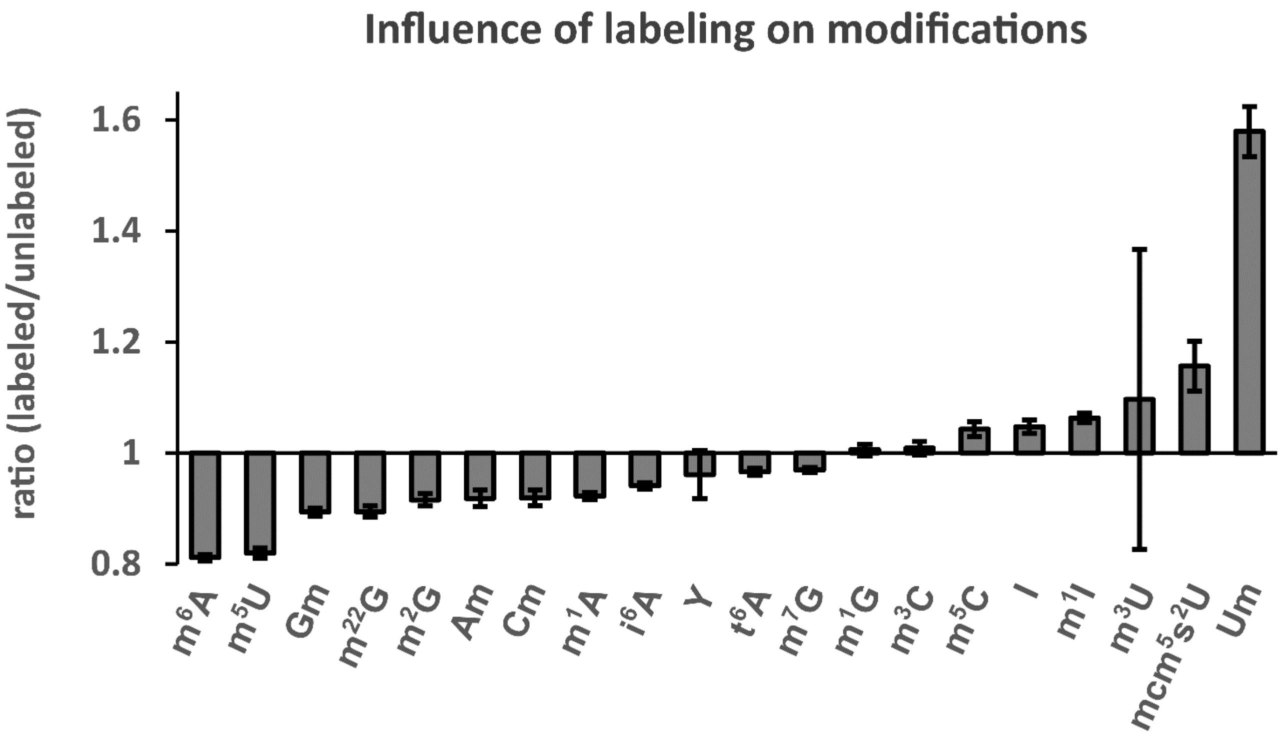


**Figure S14: Ratio of labeled to unlabeled modification amount in the mix samples.**
Cells were cultured for 7 days (=3 passages) in unlabeled or labeled media. After harvesting, the cell suspensions were mixed for subsequent co-processing. Total tRNA was purified and digested to nucleosides using standard procedures. For each modification the calculated amount of labeled modification per labeled tRNA molecule was divided by the amount of unlabeled modification per unlabeled tRNA molecule respectively. The experiment was done in n = 3 biol. replicates and error bars reflect standard deviation.

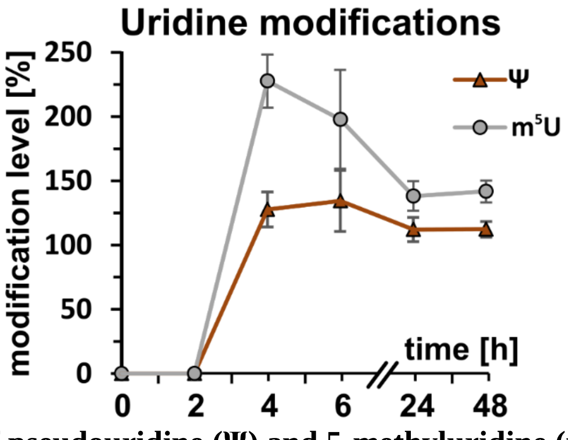


**Figure S15: Occurrence of pseudouridine (Ψ) and 5-methyluridine ($m^5U$) in new transcripts.**
Cells were grown in unlabeled DMEM D0422 (+uridine, + adenine) for 7 days. At T = 0 the medium was exchanged to DMEM D0422 supplemented with $^{15}N_5$-adenine and $^{13}C_5{}^{15}N_2$-uridine. Cells were harvested after set time points and $tRNA^{Phe}$ was purified and analyzed by LC-MS/MS. Plotted on the y-axis is the level of modification in new transcripts where 100% equals the amount of the respective nucleoside originating from unlabeled medium before experiment initiation (T = 0). The experiment was done in n = 3 biol. replicates. Error bars reflect standard deviation.

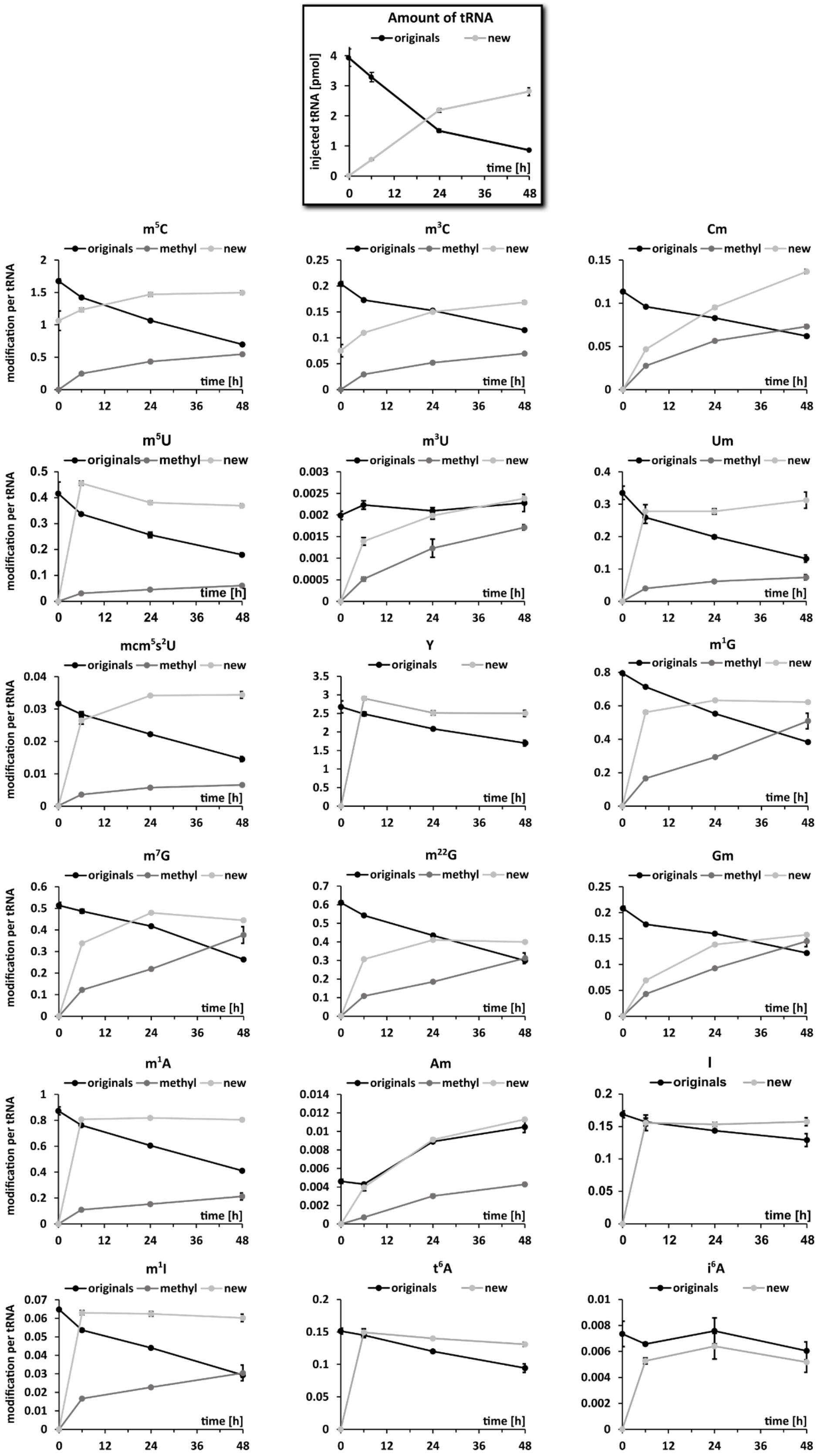


**Figure S16: Maturation processes of total tRNA in detail.**
Original nucleosides (originals, black line) already existed before experiment initiation. Post-methylated nucleosides (methyl, dark grey line) are modifications arising from the methylation of original RNA after experiment initiation. New nucleosides (new, light grey line) show the incorporation of modification into new transcripts. Data points reflect the mean and standard deviations of n = 3 biol. replicates.

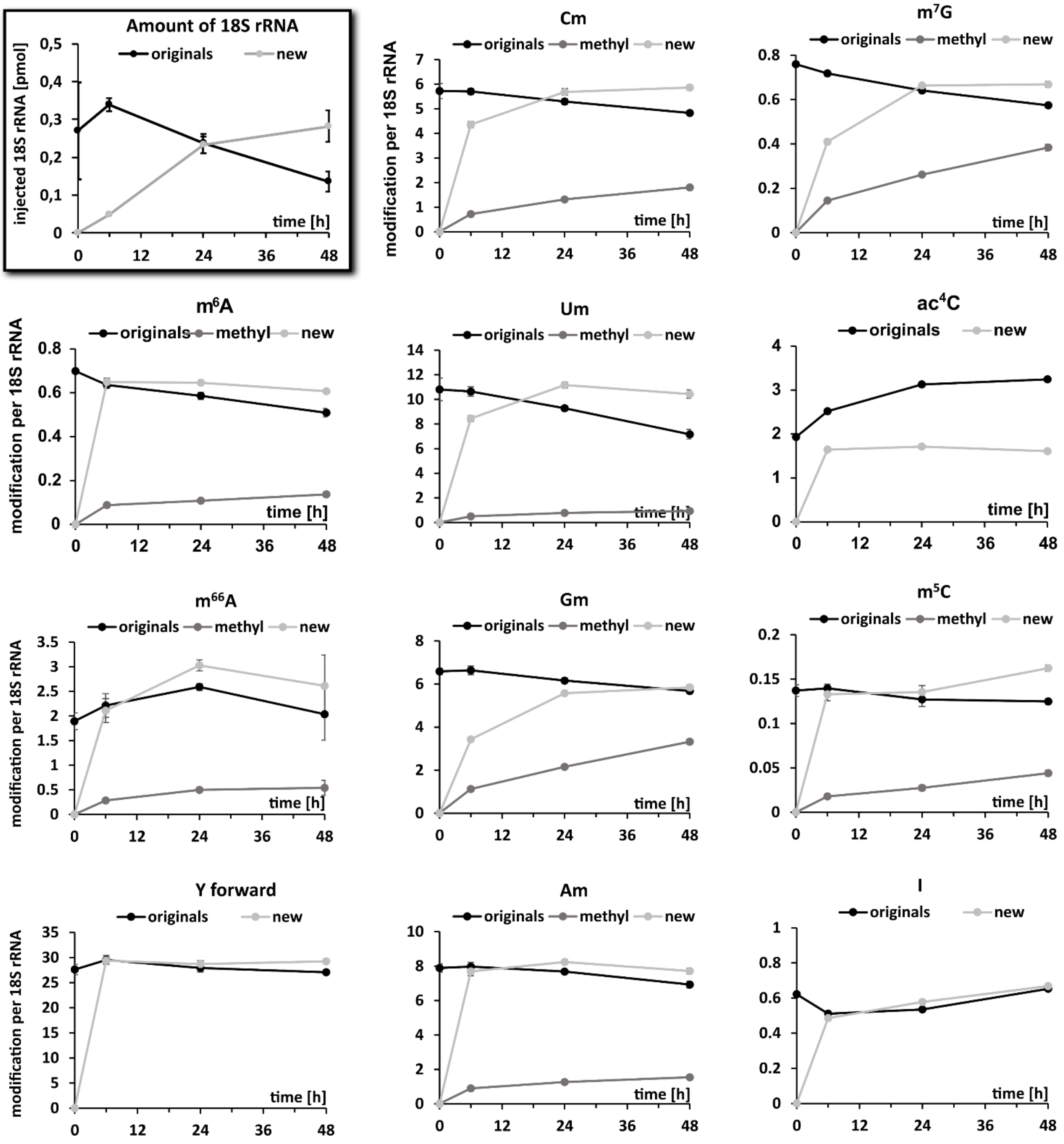


**Figure S18: Maturation processes of 18S rRNA in detail.**
Original nucleosides (originals, black line) already existed before experiment initiation. Post-methylated nucleosides (methyl, dark grey line) are modifications arising from the methylation of original RNA after experiment initiation. New nucleosides (new, light grey line) show the incorporation of modification into new transcripts. Data points reflect the mean and standard deviations of n = 3 biol. replicates.

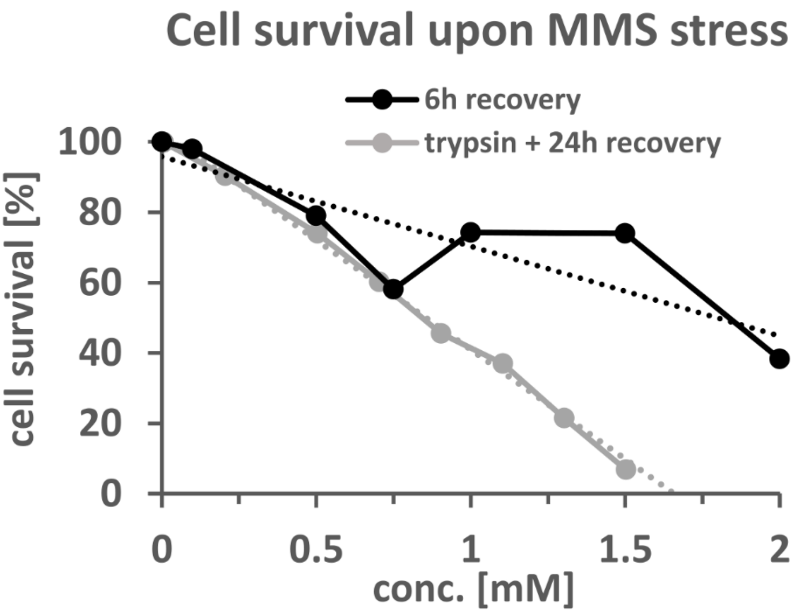


**Figure S19: Effect of MMS on growth of HEK 293 cells.**
HEK 293 cells were grown to ~70% confluency in DMEM D0422 medium supplemented with unlabeled uridine and adenine. The respective concentration of MMS was then added by exchanging the medium. After 1h the MMS containing medium was removed again and substituted by the starting medium for recovery. After 6h living cells were counted using trypan blue and a hemocytometer. For 24 h recovery the stress medium was first removed followed by trypsinization and seeding of the cells on a new plate (1:2 split). After 24 h living cells were counted using trypan blue and a hemocytometer. The dashed lines represent the respective regression curves.

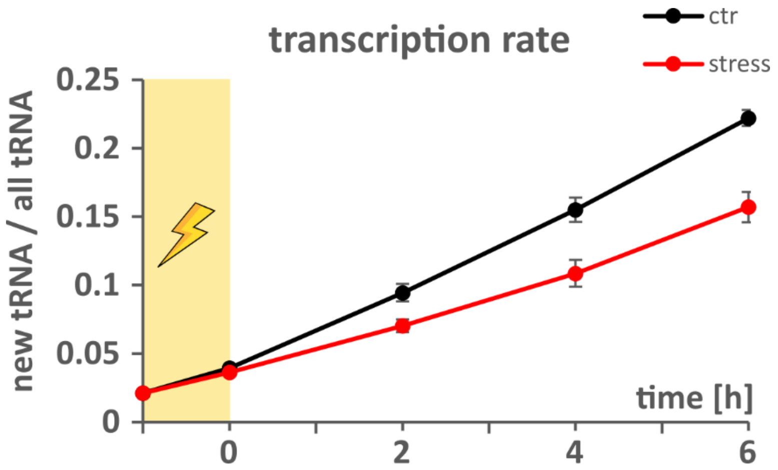


**Figure S20: Transcription rate of stressed cells.**
70% confluent $D_3$-labeled cells were incubated with fully labeled media (stable isotope labeled $^{15}N_5$-adenine, $^{13}C_5{}^{15}N_2$-uridine and $CD_3$-methionine) for 2 h before the $LD_{50}$ dose of methyl methanesulfonate (MMS, yellow shaded area) was added (T = −1). After 1 h the stress media was replaced by fresh fully labeled media. After set time points (0 h, 2 h, 4 h, 6 h) the cells were harvested and $tRNA^{Phe}_{GAA}$ was purified and subjected to LC-MS/QQQ analysis. Isoacceptor purification and digestion to nucleosides was done using standard procedures. The sum of new canonical nucleosides (labeled) was divided by the sum of all canonical nucleosides (unlabeled + labeled) and plotted against the time points for the control and stressed cells respectively. All experiments are from n = 3 biol. replicates and error bars reflect standard deviation.

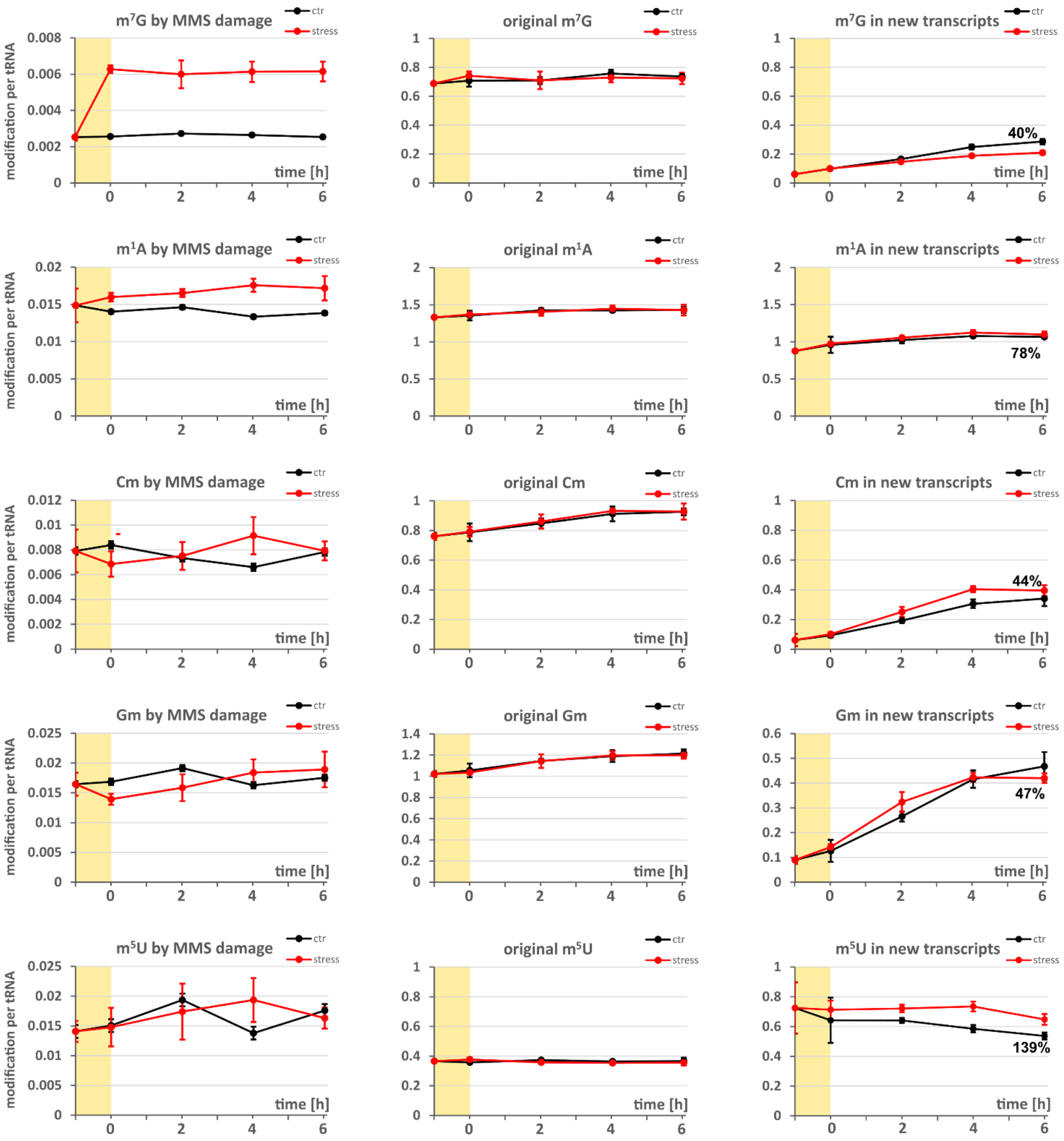
m$^7$G by MMS damage
original m$^7$G
m$^7$G in new transcripts
40%
m$^1$A by MMS damage
original m$^1$A
m$^1$A in new transcripts
78%
Cm by MMS damage
original Cm
Cm in new transcripts
44%
Gm by MMS damage
original Gm
Gm in new transcripts
47%
m$^5$U by MMS damage
original m$^5$U
m$^5$U in new transcripts
139%
ctr
stress
modification per tRNA
time [h]

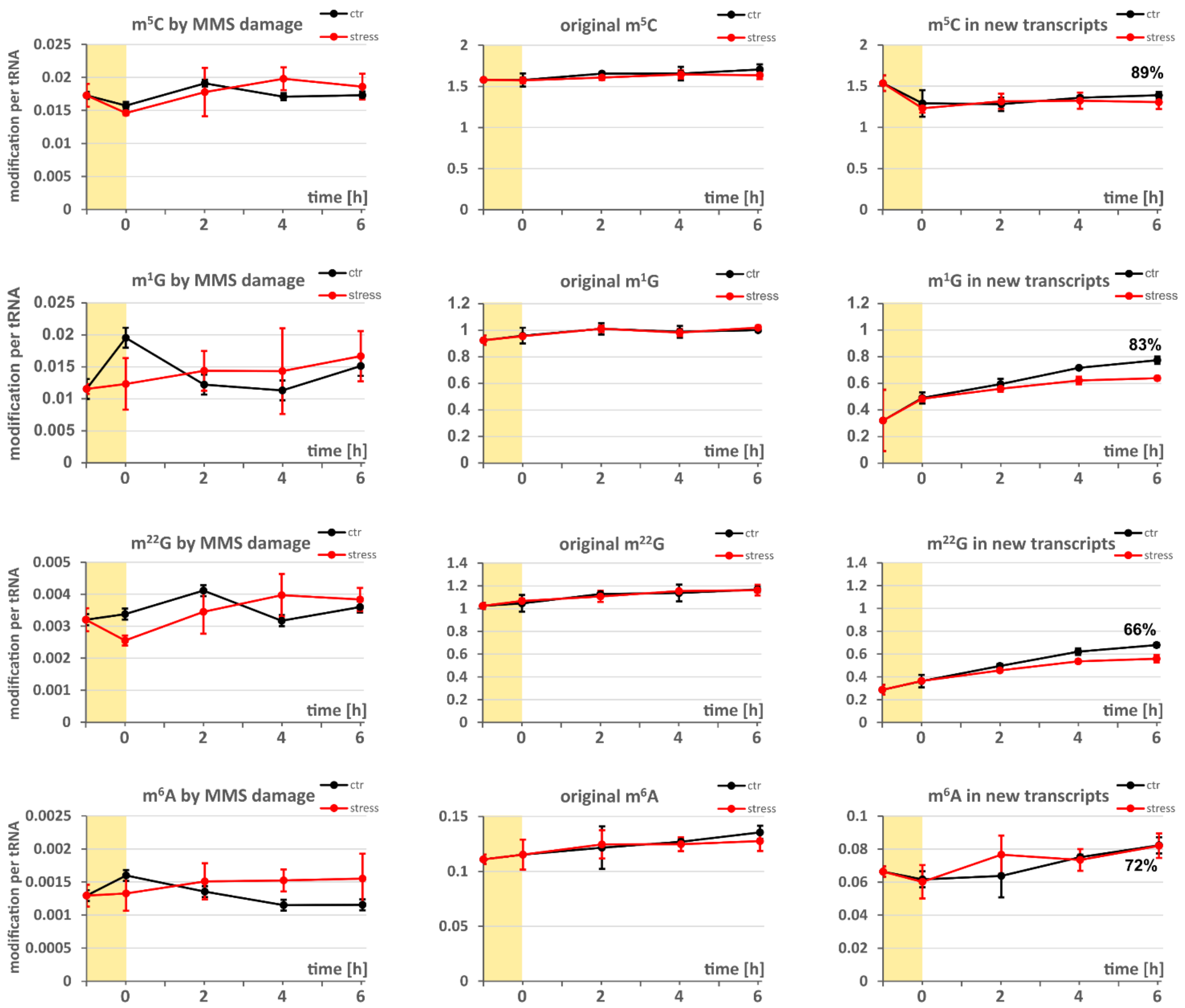


**Figure S21: Effect of MMS on tRNA modification dynamics**

70% confluent D$_3$-labeled cells were incubated with fully labeled media for 2 h before the LD$_{50}$ dose of methyl methanesulfonate (MMS, yellow shaded area) was added (T = −1). After 1 h the stress media was replaced by fresh labeled media. After set time points (0 h, 2 h, 4 h, 6 h) the cells were harvested and tRNA$^{Phe}_{GAA}$ was purified and subjected to LC-MS/QQQ analysis. Left: Modification per tRNA molecule arising from direct MMS damage in control and MMS stressed cells. Unlabeled modifications were referenced to unlabeled canonicals to calculate the amount of modifications arising from direct methylation damage by MMS. Middle: Modification per tRNA molecule in original transcripts (already existent before medium exchange at T = −1) in control and MMS stressed cells. D$_3$-labeled modifications were referenced to unlabeled canonicals to calculate the amount of modifications in original transcripts. Right: Modification per tRNA molecule in new transcripts. Labeled modifications were referenced to labeled canonicals to calculate the amount of modifications in new tRNA transcripts. The numbers at time point 6 give the percentage of modification amount in the control sample referenced to the naturally occurring amount of the respective modification (T = −1). The experiment was done in n = 3 biol. replicates and error bars reflect standard deviation

**Table S1: Results of scanned modifications for quantification of $tRNA^{Phe}$.**
The values are calculated from n = 3 biological triplicates and give the number of modifications per average $tRNA^{Phe}$ molecule. n.d., not detectable

| Modification | Average | Standard deviation |
|---|---|---|
| **Y** | 4.436 | 0.027 |
| **D** | 3.754 | 0.172 |
| **$m^1A$** | 1.464 | 0.015 |
| **$m^5C$** | 1.236 | 0.014 |
| **$m^{22}G$** | 1.042 | 0.005 |
| **Gm** | 0.990 | 0.008 |
| **Cm** | 0.909 | 0.006 |
| **$m^7G$** | 0.720 | 0.011 |
| **$m^5U$** | 0.479 | 0.011 |
| **$m^6A$** | 0.313 | 0.003 |
| **$m^1G$** | 0.278 | 0.009 |
| **I** | 0.063 | 0.003 |
| **$m^1I$** | 0.026 | 0.004 |
| **$t^6A$** | 0.016 | 0.004 |
| **$i^6A$** | 0.006 | 0.004 |
| **$m^3C$** | n.d. | |
| **$m^3U$** | n.d. | |
| **Am** | n.d. | |
| **Um** | n.d. | |
| **$mcm^5s^2U$** | n.d. | |
| **$mcm^5U$** | n.d. | |
| **$ncm^5U$** | n.d. | |
| **$m^1Y$** | n.d. | |

**Table S2: Parameters for each measured compound of MRM methods for QQQ analysis.** Nucleosides are abbreviated with the common code found at modomics [2]. “Unlabeled” refers to nucleosides from unlabeled medium; “nucleoside labeled” refers to nucleosides from $^{15}N_5$-adenine and $^{13}C_5{}^{15}N_2$-uridine medium; “methyl labeled” refers to nucleosides with no label of nucleobase or sugar but $CD_3$-methylation; “nucleoside and methyl labeled” refers to nucleosides grown in the presence of $^{15}N_5$-adenine, $^{13}C_5{}^{15}N_2$-uridine and $CD_3$-methionine; “SILIS” stands for stable isotope labeled internal standard and was produced in yeast using a rich $^{15}N/^{13}C$ growth medium following our published procedure [3].

| | Compound Name | Precursor Ion | Product Ion | Ret Time (min) | Fragmentor (V) | Collision Energy (eV) |
|---|---|---|---|---|---|---|
| unlabeled | A | 268 | 136 | 5,2 | 200 | 20 |
| | acp$^{3}$U | 346 | 214 | 2,3 | 95 | 15 |
| | Am | 282 | 136 | 6,0 | 130 | 17 |
| | C | 244 | 112 | 2,1 | 200 | 20 |
| | Cm | 258 | 112 | 4,1 | 180 | 9 |
| | D | 247 | 115 | 1,6 | 70 | 5 |
| | G | 284 | 152 | 4,3 | 200 | 20 |
| | Gm | 298 | 152 | 5,0 | 100 | 9 |
| | I | 269 | 137 | 4,1 | 100 | 10 |
| | i$^{6}$A | 336 | 204 | 8,0 | 140 | 17 |
| | m$^{1}$A | 282 | 150 | 2,5 | 150 | 25 |
| | m$^{1}$G | 298 | 166 | 4,9 | 105 | 13 |
| | m$^{1}$I | 283 | 151 | 4,8 | 80 | 12 |
| | m$^{1}$Y | 259 | 223 | 3,1 | 85 | 5 |
| | m$^{22}$G | 312 | 180 | 5,7 | 105 | 13 |
| | m$^{2}$G | 298 | 166 | 5,1 | 95 | 17 |
| | m$^{3}$C | 258 | 126 | 2,3 | 88 | 14 |
| | m$^{3}$U | 259 | 127 | 4,8 | 75 | 9 |
| | m$^{5}$C | 258 | 126 | 3,8 | 185 | 13 |
| | m$^{5}$U | 259 | 127 | 4,4 | 95 | 9 |
| | m$^{6}$A | 282 | 150 | 6,5 | 125 | 17 |
| | m$^{7}$G | 298 | 166 | 3,6 | 100 | 13 |
| | mcm$^{5}$s$^{2}$U | 333 | 201 | 6,2 | 92 | 8 |
| | t$^{6}$A | 413 | 281 | 5,8 | 130 | 9 |
| | U | 245 | 113 | 3,0 | 95 | 5 |
| | Um | 259 | 113 | 4,6 | 96 | 8 |
| | Y | 245 | 209 | 1,7 | 90 | 5 |
| | mcm$^{5}$U | 317 | 185 | 5,0 | 95 | 5 |
| | ncm$^{5}$U | 302 | 170 | 2,5 | 85 | 8 |
| nucleos | A lab | 273 | 141 | 5,2 | 200 | 20 |

| | Compound Name | Precursor Ion | Product Ion | Ret Time (min) | Fragmentor (V) | Collision Energy (eV) |
|---|---|---|---|---|---|---|
| | acp$^3$U lab | 353 | 216 | 2,3 | 95 | 15 |
| | Am lab | 287 | 141 | 6,0 | 130 | 17 |
| | C lab | 251 | 114 | 2,1 | 200 | 20 |
| | Cm lab | 265 | 114 | 4,1 | 180 | 9 |
| | D lab | 254 | 117 | 1,6 | 70 | 5 |
| | G lab | 288 | 156 | 4,3 | 200 | 20 |
| | Gm lab | 302 | 156 | 5,0 | 100 | 9 |
| | I lab | 273 | 141 | 4,1 | 100 | 10 |
| | i$^6$A lab | 341 | 209 | 8,0 | 140 | 17 |
| | m$^1$A lab | 287 | 155 | 2,5 | 150 | 25 |
| | m$^1$G lab | 302 | 170 | 4,9 | 105 | 13 |
| | m$^1$I lab | 287 | 155 | 4,8 | 80 | 12 |
| | m$^1$Y lab | 266 | 230 | 3,1 | 85 | 5 |
| | m$^{22}$G lab | 316 | 184 | 5,7 | 105 | 13 |
| | m$^2$G lab | 302 | 170 | 5,1 | 95 | 17 |
| | m$^3$C lab | 265 | 128 | 2,3 | 88 | 14 |
| | m$^3$U lab | 266 | 129 | 4,8 | 75 | 9 |
| | m$^5$C lab | 265 | 128 | 3,8 | 185 | 13 |
| | m$^5$U lab | 266 | 129 | 4,4 | 95 | 9 |
| | m$^6$A lab | 287 | 155 | 6,5 | 125 | 17 |
| | m$^7$G lab | 302 | 170 | 3,6 | 100 | 13 |
| | mcm$^5$s$^2$U lab | 340 | 203 | 6,2 | 92 | 8 |
| | t$^6$A lab | 418 | 286 | 5,8 | 130 | 9 |
| | U lab | 252 | 115 | 3,0 | 95 | 5 |
| | Um lab | 266 | 115 | 4,6 | 96 | 8 |
| | Y lab | 252 | 216 | 1,7 | 90 | 5 |
| | mcm$^5$U lab | 324 | 187 | 5,0 | 95 | 5 |
| | ncm$^5$U lab | 309 | 172 | 2,5 | 85 | 8 |
| methyl labeled | Am $D_3$ | 285 | 136 | 6,0 | 130 | 17 |
| | Cm $D_3$ | 261 | 112 | 4,1 | 180 | 9 |
| | Gm $D_3$ | 301 | 152 | 5,0 | 100 | 9 |
| | m$^1$A $D_3$ | 285 | 153 | 2,5 | 150 | 25 |
| | m$^1$G $D_3$ | 301 | 169 | 4,9 | 105 | 13 |
| | m$^1$I $D_3$ | 286 | 154 | 4,8 | 80 | 12 |
| | m$^1$Y $D_3$ | 262 | 226 | 3,1 | 85 | 5 |
| | m$^{22}$G $D_3$ | 318 | 186 | 5,7 | 105 | 13 |
| | m$^2$G $D_3$ | 301 | 169 | 5,1 | 95 | 17 |
| | m$^3$C $D_3$ | 261 | 129 | 2,3 | 88 | 14 |
| | m$^3$U $D_3$ | 262 | 130 | 4,8 | 75 | 9 |

| | Compound Name | Precursor Ion | Product Ion | Ret Time (min) | Fragmentor (V) | Collision Energy (eV) |
|---|---|---|---|---|---|---|
| | $m^5C\ D_3$ | 261 | 129 | 3,8 | 185 | 13 |
| | $m^5U\ D_3$ | 262 | 130 | 4,4 | 95 | 9 |
| | $m^6A\ D_3$ | 285 | 153 | 6,5 | 125 | 17 |
| | $m^7G\ D_3$ | 301 | 169 | 3,6 | 100 | 13 |
| | $mcm^5s^2U\ D_3$ | 336 | 204 | 6,2 | 92 | 8 |
| | $Um\ D_3$ | 262 | 113 | 4,6 | 96 | 8 |
| | $mcm^5U\ D_3$ | 320 | 188 | 5,0 | 95 | 5 |
| nucleoside and methyl labeled | $Am\ lab\ D_3$ | 290 | 141 | 6,0 | 130 | 17 |
| | $Cm\ lab\ D_3$ | 268 | 114 | 4,1 | 180 | 9 |
| | $Gm\ lab\ D_3$ | 305 | 156 | 5,0 | 100 | 9 |
| | $m^1A\ lab\ D_3$ | 290 | 158 | 2,5 | 150 | 25 |
| | $m^1G\ lab\ D_3$ | 305 | 173 | 4,9 | 105 | 13 |
| | $m^1I\ lab\ D_3$ | 290 | 158 | 4,8 | 80 | 12 |
| | $m^1Y\ lab\ D_3$ | 269 | 233 | 3,1 | 85 | 5 |
| | $m^{22}G\ lab\ D_3$ | 322 | 190 | 5,7 | 105 | 13 |
| | $m^2G\ lab\ D_3$ | 305 | 173 | 5,1 | 95 | 17 |
| | $m^3C\ lab\ D_3$ | 268 | 131 | 2,3 | 88 | 14 |
| | $m^3U\ lab\ D_3$ | 269 | 132 | 4,8 | 75 | 9 |
| | $m^5C\ lab\ D_3$ | 268 | 131 | 3,8 | 185 | 13 |
| | $m^5U\ lab\ D_3$ | 269 | 132 | 4,4 | 95 | 9 |
| | $m^6A\ lab\ D_3$ | 290 | 158 | 6,5 | 125 | 17 |
| | $m^7G\ lab\ D_3$ | 305 | 173 | 3,6 | 100 | 13 |
| | $mcm^5s^2U\ lab\ D_3$ | 343 | 206 | 6,2 | 92 | 8 |
| | $Um\ lab\ D_3$ | 269 | 115 | 4,6 | 96 | 8 |
| | $mcm^5U\ lab\ D_3$ | 327 | 190 | 5,0 | 95 | 5 |
| SILIS | A SILIS | 283 | 146 | 5,2 | 200 | 20 |
| | Am SILIS | 298 | 146 | 6,0 | 130 | 17 |
| | C SILIS | 256 | 119 | 2,1 | 200 | 20 |
| | Cm SILIS | 271 | 119 | 4,1 | 180 | 9 |
| | D SILIS | 258 | 121 | 1,6 | 70 | 5 |
| | G SILIS | 299 | 162 | 4,3 | 200 | 20 |
| | Gm SILIS | 314 | 162 | 5,0 | 100 | 9 |
| | I SILIS | 283 | 146 | 4,1 | 100 | 10 |
| | $i^6A$ SILIS | 356 | 219 | 8,0 | 140 | 17 |
| | $m^1A$ SILIS | 298 | 161 | 2,5 | 150 | 25 |
| | $m^1G$ SILIS | 314 | 177 | 4,9 | 105 | 13 |
| | $m^1I$ SILIS | 298 | 161 | 4,8 | 80 | 12 |

| | Compound Name | Precursor Ion | Product Ion | Ret Time (min) | Fragmentor (V) | Collision Energy (eV) |
|---|---|---|---|---|---|---|
| | m$^{22}$G SILIS | 329 | 192 | 5,7 | 105 | 13 |
| | m$^{2}$G SILIS | 314 | 177 | 5,1 | 95 | 17 |
| | m$^{3}$C SILIS | 271 | 134 | 2,3 | 88 | 14 |
| | m$^{5}$C SILIS | 271 | 134 | 3,8 | 185 | 13 |
| | m$^{5}$U SILIS | 271 | 134 | 4,4 | 95 | 9 |
| | m$^{6}$A SILIS | 298 | 161 | 6,5 | 125 | 17 |
| | m$^{7}$G SILIS | 314 | 177 | 3,6 | 100 | 13 |
| | mcm$^{5}$s$^{2}$U SILIS | 347 | 210 | 6,2 | 92 | 8 |
| | t$^{6}$A SILIS | 434 | 297 | 5,8 | 130 | 9 |
| | U SILIS | 256 | 119 | 3,0 | 95 | 5 |
| | Um SILIS | 271 | 119 | 4,6 | 96 | 8 |
| | Y SILIS | 256 | 220 | 1,7 | 90 | 5 |
| | mcm$^{5}$U SILIS | 331 | 194 | 5,0 | 95 | 5 |
| | ncm$^{5}$U SILIS | 316 | 179 | 2,5 | 85 | 8 |
